# F-actin dynamics couples sphingolipid metabolism to epithelial barrier integrity in chronic colitis

**DOI:** 10.64898/2026.01.21.700747

**Authors:** Snezhanna Medvedeva, Julia Popova, Kseniya Achasova, Julia Kulygina, Ekaterina Nickelwart, Lyubov Suldina, Ksenia Morozova, Marina Osipenko, Elena Kozhevnikova

## Abstract

**Background:** Intestinal barrier dysfunction is a hallmark of inflammatory bowel diseases (IBD). This condition causes intoxication and immune hyperactivation. Understanding the events underlying epithelial barrier disruption during chronic inflammation is key to developing barrier-restoring therapies. Filamentous actin (F-actin) is essential for maintaining polarity and junctional integrity. However, the contribution of F-actin dynamics to IBD-associated barrier dysfunction remains unclear.

**Objective:** We aimed to examine actin cytoskeleton integrity during chronic colitis across mouse models and human patients and identify potential regulators of cytoskeleton dynamics.

**Design:** F-actin and junctional proteins were analyzed in three models of chronic colitis (*Muc2* KO, DSS-induced colitis, adoptive transfer colitis) using confocal microscopy. Claudin-3 interactors were identified by immunoprecipitation and proteomics. Intestinal organoids were used to assess the effect of F-actin disruption on barrier integrity. Metabolomic and gene expression analyses identified candidate pathways, further validated by chemical inhibition. Biopsies from patients with ulcerative colitis (UC) were examined using transmission electron microscopy and confocal microscopy.

**Results:** Disrupted actin dynamics emerged as a critical driver of epithelial barrier dysfunction in chronic colitis. An imbalance between polymeric and monomeric actin impaired barrier integrity *in vivo* and in 3D organoids. Immunoprecipitation identified actin and associated factors as the primary interactors of claudin-3 with reduced interaction during inflammation. Ceramide metabolism was revealed as a potential regulator of F-actin and intestinal barrier. In UC patients, we confirmed concurrent disruption of junctions and F-actin.

**Conclusions:** F-actin dysregulation contributes to barrier dysfunction in IBD and targeting its modulators, including ceramide biosynthesis, represents a novel therapeutic strategy.

**WHAT IS ALREADY KNOWN ON THIS TOPIC:**

- Epithelial damage and increased paracellular permeability are key characteristics of inflammatory bowel diseases.
- Paracellular permeability is partially attributed to the downregulation of junction proteins but this mechanism does not explain all clinical observations.
- In the *Muc2* KO mouse model of chronic colitis, F-actin organization and membrane localization of tight junction protein claudin-3 are disrupted, although protein expression levels remain unchanged.

**WHAT THIS STUDY ADDS:**

- F-actin dynamics is impaired in the intestinal epithelium across three different mouse models of chronic colitis and IBD patients.
- Disruption of F-actin dynamics leads to impaired membrane localization of tight and adherens junction proteins and increased intestinal epithelial permeability *in vivo* and in colonic organoids.
- Inhibition of ceramide biosynthesis rescues F-actin polymerization and intestinal barrier integrity in mouse chronic colitis models.

**HOW THIS STUDY MIGHT AFFECT RESEARCH, PRACTICE OR POLICY:**

- Targeting F-actin dynamics is a promising approach to improve gut epithelial integrity.
- "Ceramide–F-actin–junction" axis is proposed as one of the mechanisms behind epithelial barrier disruption in colitis. Therapeutic targeting of this axis represents a promising path for restoring gut integrity.

## Introduction

The intestinal epithelium forms a selective semi-permeable barrier which isolates luminal microbiota, pathogens, parasites, and food particles from the internal environment. Numerous disorders are associated with increased intestinal permeability including inflammatory bowel diseases (IBD), type 1 diabetes mellitus, celiac disease, multiple sclerosis etc. Disrupted intestinal barrier facilitates the invasion of pathogens into the lamina propria and exaggerates the mucosal immune response (1,2). Mucus deficiency contributes to the initiation and persistence of chronic inflammation in IBD (3)

IBD are chronic, relapsing idiopathic disorders of the gastrointestinal tract. This term encompasses two major conditions that differ in anatomical distribution and depth of inflammation. Ulcerative colitis (UC) is characterized by continuous inflammation in the colonic mucosa, most commonly affecting the rectum, but potentially extending proximally throughout the entire colon. In contrast, Crohn’s disease (CD) causes transmural inflammation that can affect any region of the gastrointestinal tract, preferentially involving the terminal ileum and colon (4).

As a chronic relapsing condition, IBD is characterized by periods of active and inactive inflammation. Microscopic evaluation of patients with active IBD reveals pronounced infiltration of immune cells into the lamina propria, cryptitis, granuloma formation. Epithelial structure is abruptly changed during active IBD. Biopsy specimens show tissue damage: mucosal ulcers, crypt atrophy, and abscesses (5). Inactive IBD refers to the remission state, which can be subdivided into clinical (no symptoms), endoscopic (mucosal healing), histological (normal epithelium), and biochemical (normal levels of inflammatory markers) remissions (6,7). These kinds of remissions do not always correlate: mucosal healing may occur in the absence of clinical remission; microscopic features of inflammation may persist in clinical and endoscopic remission (8–10). Increased permeability and epithelium leakage persist during inactive IBD, establishing predisposition for recurrent inflammation. The mechanisms underlying persistent epithelial barrier dysfunction in quiescent IBD are poorly understood, despite this state of low-grade inflammation and increased permeability being the most common clinical manifestation.

The protective intestinal barrier is primarily formed by apical junctional complexes: tight junctions (TJs), adherens junctions (AJs) and desmosomes, which connect epithelial cells and regulate selective paracellular transport of ions, water, and small molecules while preventing translocation of luminal antigens and pathogens into the blood (1,11,12). TJs and AJs are associated with a network of cortical actin microfilaments underlying the plasma membrane. Filamentous actin (F-actin) also anchors epithelial cells to the basement membrane via focal adhesions and forms the structural core of apical microvilli (13). Normally, dynamic remodeling of junctions and the actin cytoskeleton facilitates immune cell trafficking across the epithelium (14). In chronic inflammation, dysregulation of these processes disrupts mucosal homeostasis, leading to a pathologic feedback loop that perpetuates tissue damage and inflammation.

The expression of TJ proteins, including claudins, ZO-1, and occludin, is altered in active colitis (15,16). Similar alterations were observed in AJ proteins, specifically E-cadherin and β-catenin (17,18). Data on TJ and AJ protein expression during inactive colitis are limited, although some patient studies demonstrate no significant alterations (16). Notably, the existing *in vivo* data on F-actin rearrangements during inflammation are both limited and contradictory (19,20). Therefore, the molecular crosstalk between actin microfilaments and junctional complexes is emerging as a critical determinant of intestinal barrier permeability.

Numerous IBD animal models enable investigating various aspects of IBD. These include chemically- and immunologically-induced, spontaneous, genetically-engineered, and T-cell transfer models (21,22). DSS administration can be adapted to model either acute or chronic colitis, whereas KO models such as *Muc2^-/-^* mice, spontaneously develop chronic intestinal inflammation (21,23). T-cell-dependent models of IBD are significantly more relevant to human chronic disease, given the central role of T-cells in the disease pathogenesis (24).

Our previous work in the *Muc2* KO model revealed key features of chronic inflammation, namely F-actin rearrangement and claudin-3 delocalization from the lateral membrane (25). Given these data, we hypothesized that F-actin instability in the enterocytes is a key driver of TJ and AJ disassembly. Here, we demonstrate that dysregulated actin cytoskeleton dynamics represent a conserved pathological feature of chronic intestinal inflammation, shared across three independent animal models and UC patients. The functional link between F-actin disruption and increased intestinal permeability is supported by *in vivo* and *in vitro* functional assays, as well as the analysis of the claudin-3 protein complex. We further identified ceramide as a potent metabolic regulator of F-actin dynamics and barrier function upon chronic colitis.

## Results

### Claudin-7, β-catenin, and E-cadherin but not ZO-1 are delocalized from the lateral membranes of enterocytes in *Muc2* KO mice

We analyzed the subcellular distribution of TJ and AJ proteins in the descending colon of *Muc2* KO mice to assess epithelial barrier defects. In previous work, we demonstrated a reduction of F-actin and the mislocalization of the tight junction protein claudin-3 (25). Neither claudin-3 nor β-actin were downregulated at the protein level, indicating that the observed differences cannot be explained by the changes in gene expression. Likewise, *Cldn7* and *ZO-1* gene expression remained unchanged in colonic samples of *Muc2* KO mice (25). In the present study, we further investigated protein localization by confocal microscopy to a broader panel of junctional proteins to provide a more comprehensive view of barrier defects.

Immunofluorescence analysis revealed that TJ protein claudin-7, but not ZO-1, is delocalized from the lateral membrane of the *Muc2* KO colonocytes (Figure 1 A, B) (*Muc2^-/-^* vs *Muc2^+/+^*, Mann-Whitney *U*-test, claudin-7: Z = -3.49, p_adj_ < 0.001; JAM-A: Z = 2.39, p_adj_ < 0.05; ZO-1: Z = 1.35, p_adj_ > 0.05). Likewise, AJ proteins E-cadherin and β-catenin exhibited weak localization to the plasma membrane, as confirmed by fluorescence intensity quantification (Figure 1 A, B) (*Muc2^-/-^* vs *Muc2^+/+^*, Mann-Whitney *U*-test, E-cadherin: Z = -4.43, p_adj_ < 0.001; β-catenin: Z = -7.05, p_adj_ < 0.001). According to the Western blot analysis, the expression of E-cadherin and β-catenin was not altered in *Muc2* KO mice similar to that earlier shown for claudins (Figure 1C) (*Muc2^-/-^* vs *Muc2^+/+^*, Mann-Whitney *U*-test, E-cadherin: Z = 0.29, p > 0.05; β-catenin: Z = -1.14, p > 0.05) (25). Consistent with our previous study, both apical and lateral F-actin were reduced (25). Thus, chronic inflammation destabilizes the actin cytoskeleton and causes mislocalization of multiple junctional proteins, including claudin-7, E-cadherin, and β-catenin, in addition to the previously published claudin-3, leading to intestinal barrier disruption. For further experiments, claudin-7 and β-catenin were used as TJ and AJ markers, respectively.

**Figure 1.**
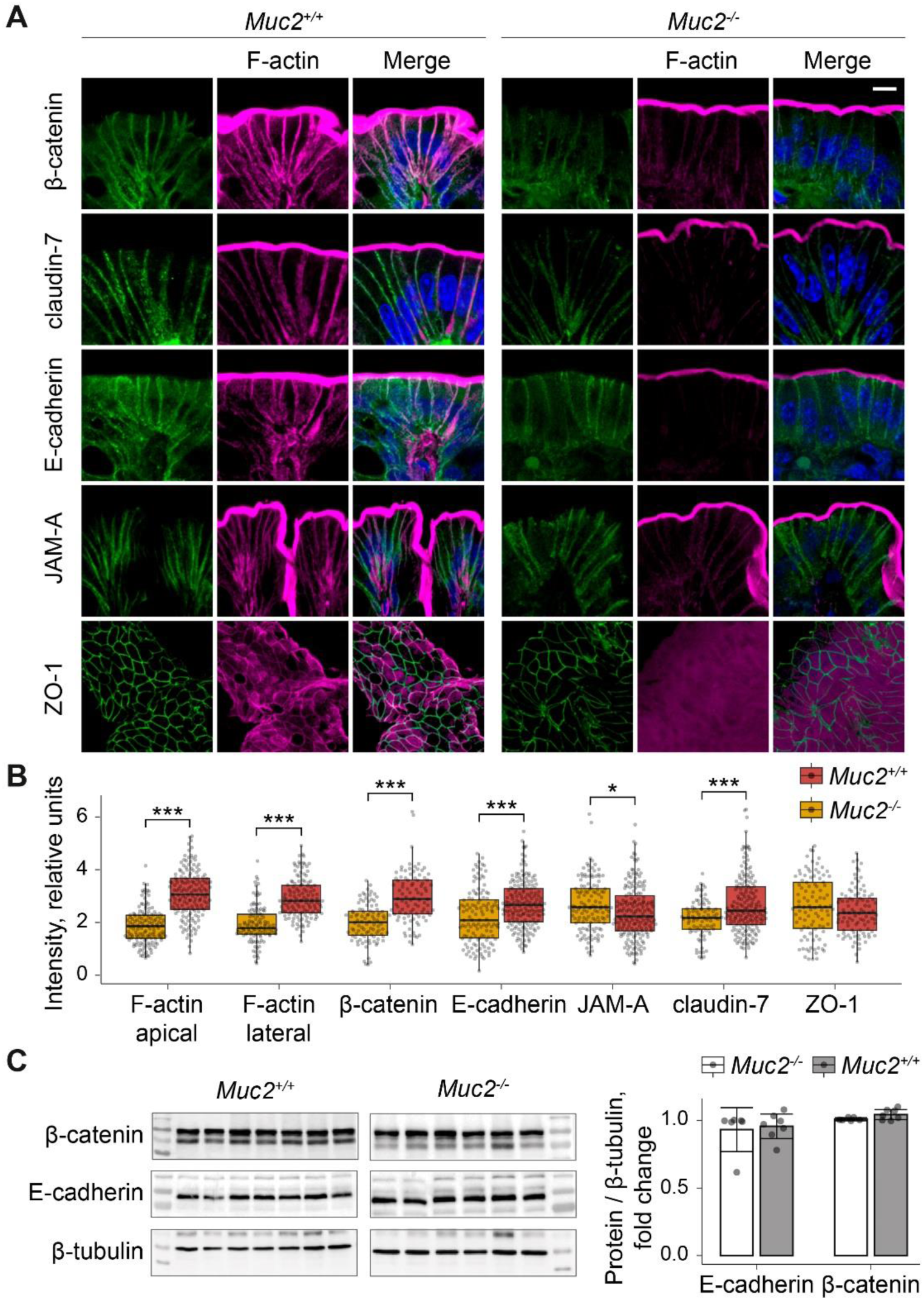
Membrane localization of adherens junction (AJ) and tight junction (TJ) proteins is compromised in the *Muc2* KO model of chronic colitis. (A) Immunostaining of the AJ and TJ proteins in the descending colon of *Muc2* KO mice. ZO-1 is shown as the view from the lumen. Bar ‒ 10 µm. (B) Fluorescence intensity quantification of AJ and TJ proteins along the cell membrane (\**p* < 0.05, \*\*\**p* < 0.001, n = 3, Mann-Whitney *U*-test). (C) Western blot analysis of the total protein in colonic samples. β-tubulin (55 kDa) was used as a loading control. Data are shown as mean ± 95% confidence interval.

### F-actin dynamics is essential for proper epithelial barrier formation in *Muc2* KO model of chronic colitis

Using claudin-3-specific antibodies for co-immunoprecipitation followed by LC-MS/MS, we identified proteins associated with claudin-3 in colonic tissues of *Muc2* KO mice and their wild-type littermates (Figure 2). In control tissues, claudin-3 co-purified primarily with a network of actin-binding proteins, including Arp2/3, α-actinin, ezrin, vinculin, actin and others (Figure 2). This specific interactome was largely lost in *Muc2* KO animals. Importantly, the total levels of the recovered claudin-3 protein were similar in both groups, as confirmed by identical intensity-based absolute quantification (iBAQ) scores, indicating post-translational disruption of its regulatory complex. These data demonstrate that intestinal barrier dysfunction in *Muc2* KO mice is closely connected to the actin protein interaction network.

**Figure 2.**
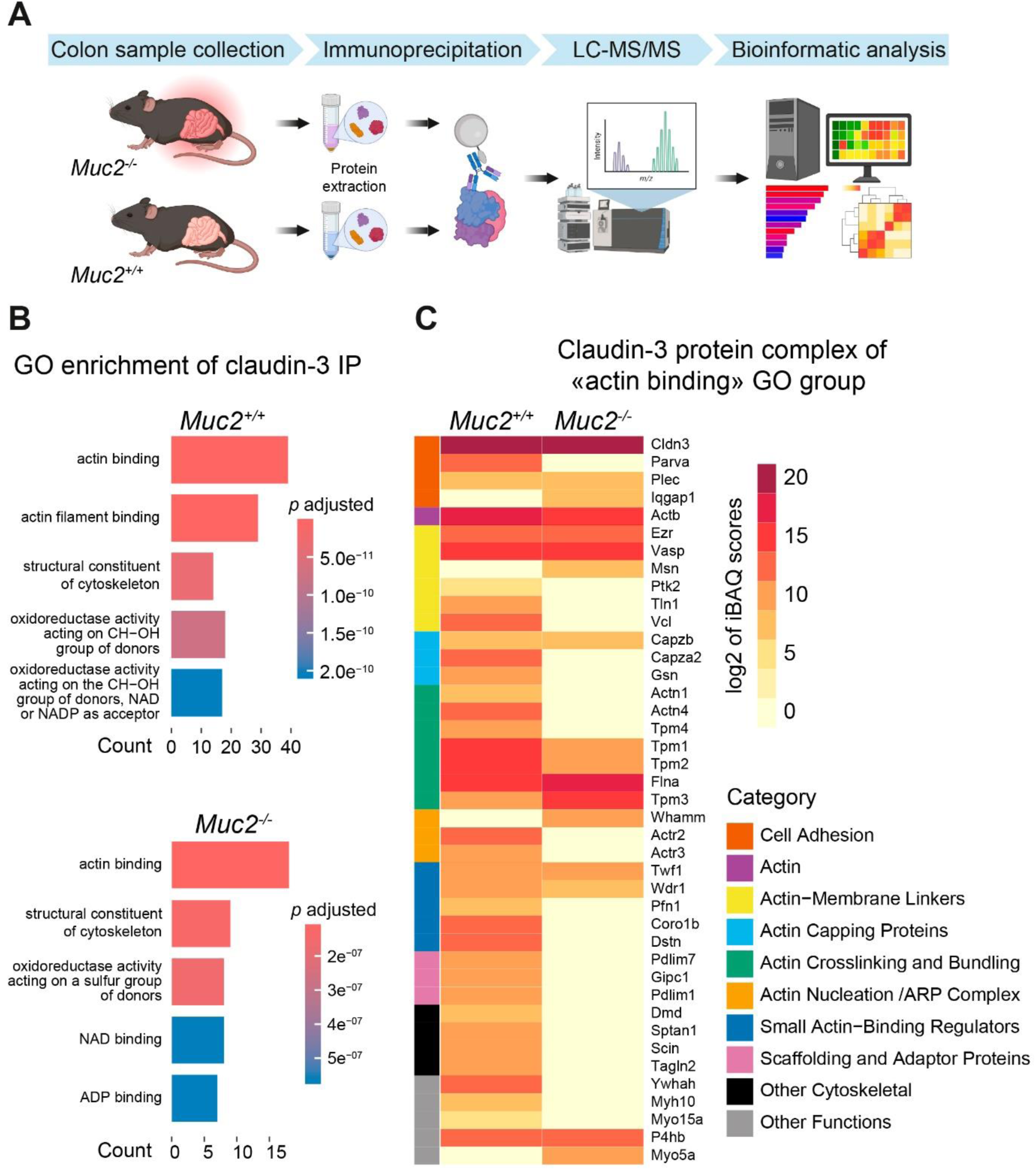
Comparative analysis of the claudin-3 interactome in *Muc2^+/+^* and *Muc2* KO mice. (A) Interactome analysis flowchart. (B) Gene Ontology enrichment of claudin-3 interactome in *Muc2* KO mice. (C) Heatmap of log₂ iBAQ abundance scores for proteins annotated with Gene Ontology “actin binding” group detected in claudin-3 immunoprecipitants from colon tissue of *Muc2* KO mice and their wild-type littermates.

Given that at least claudin-3, one of the major intestinal junctional components, interacts mostly with actin and its binding network, rather than other TJ proteins, F-actin disruption is likely to impact TJ structure. To determine whether junctional protein mislocalization is linked to F-actin disruption, we pharmacologically altered actin dynamics *in vivo* using actin polymerization drugs latrunculin A (LatA) and jasplakinolide (JP), or the ROCK inhibitor Y-27632, administered rectally to wild-type C57BL/6 mice. Immunofluorescence analysis revealed that all three treatments significantly compromised apical and lateral F-actin architecture, despite the opposing mechanisms of LatA and JP (Figure 3A-F, F-actin) (Control vs LatA, Mann-Whitney *U*-test, apical F-actin: Z = 8.56, p_adj_ < 0.001; lateral F-actin: Z = 11.96, p_adj_ < 0.001; Control vs JP, Mann-Whitney *U*-test, apical F-actin: Z = 8.64, p_adj_ < 0.001; lateral F-actin: Z = 6.75, p_adj_ < 0.001; Control vs Y-27623, Mann-Whitney *U*-test, lateral F-actin: Z = 2.26, p_adj_ < 0.05). This indicates that cytoskeletal integrity depends on the intricate balance between G-actin and F-actin. Shifting this balance towards either direction compromises cytoskeletal integrity.

**Figure 3.**
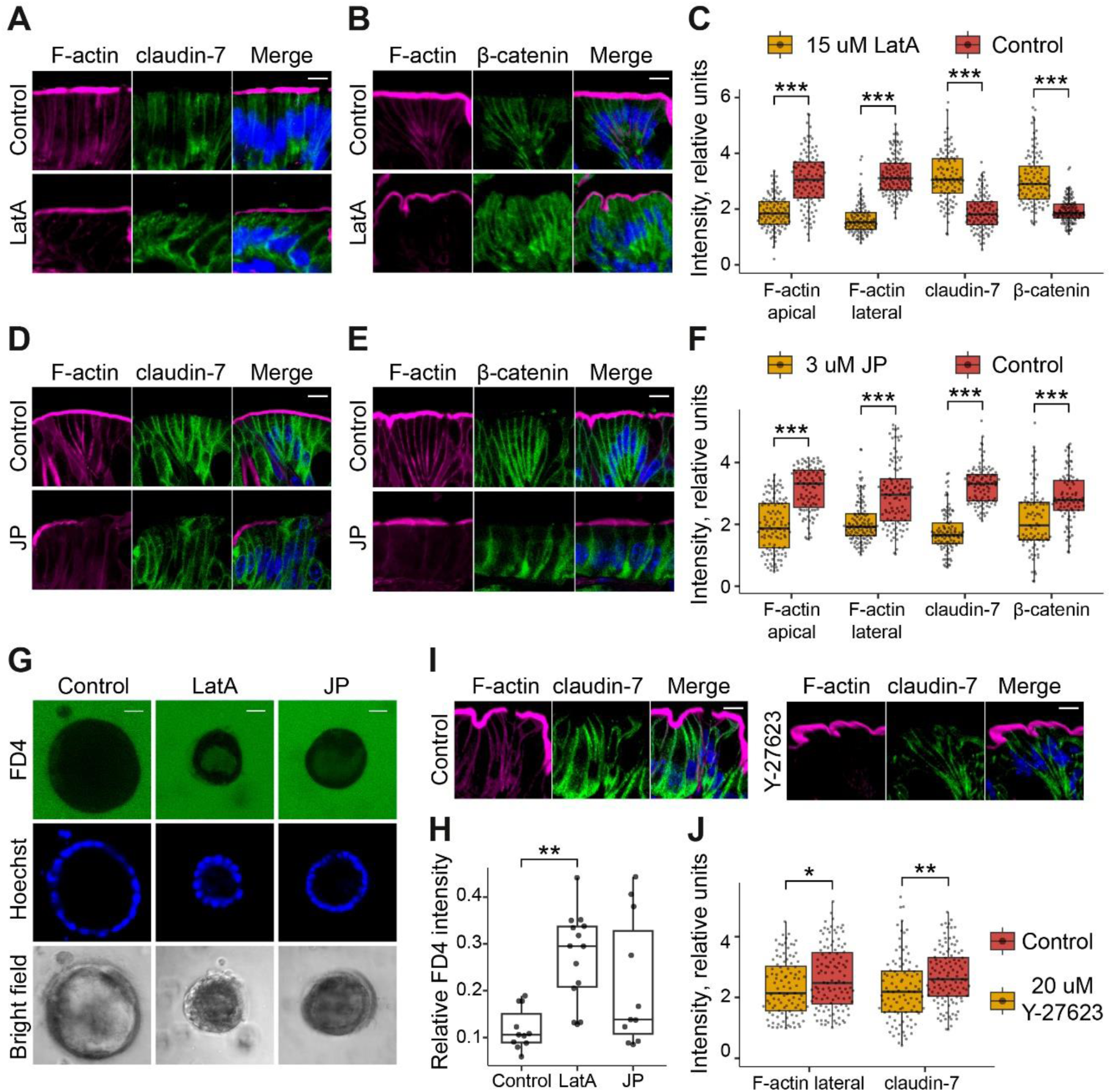
Actin dynamics controls membrane localization of AJ and TJ proteins, as well as epithelial permeability. (A) Claudin-7 and (B) β-catenin immunostaining in the descending colon of C57BL/6 mice after rectal administration of latrunculin A (LatA) for 90 min. (C) Fluorescence intensity quantification of F-actin, claudin-7 and along the cell membrane of C57BL/6 gut treated by latrunculin A (\*\*\**p* < 0.001, n = 3, Mann-Whitney *U*-test). (D) Claudin-7 and (E) β-catenin immunostaining in the descending colon of C57BL/6 mice after rectal administration of jasplakinolide (JP) for 90 min. (F) Fluorescence intensity quantification of F-actin, claudin-7 and β-catenin along the cell membrane of C57BL/6 gut treated by jasplakinolide (\*\*\**p* < 0.001, n = 3, Mann-Whitney *U*-test). (G) Latrunculin A and jasplakinolide cause permeability of the epithelial barrier in mouse intestinal organoids. Bar ‒ 10 µm. (H) 4 kDa FITC-Dextran (FD4) fluorescence relative intensity quantification in organoids (\*\**p* < 0.01, one-way ANOVA with Tukey’s post-hoc test). (I) Claudin-7 immunostaining in the descending colon of C57BL/6 mice after rectal administration of Y-27623 for 9 hours. (J) Fluorescence intensity quantification of F-actin and claudin-7 along the cell membrane of C57BL/6 gut treated by Y-27623 for 9 hours (\*\*\**p* < 0.001, n = 3, Mann-Whitney *U*-test).

Actin cytoskeleton rearrangement determined the localization of the junctional proteins claudin-7 and β-catenin (Figure 3 A-F) (Control vs LatA, Mann-Whitney *U*-test, claudin-7: Z = - 9.29, p_adj_ < 0.001; β-catenin: Z = -9.9, p_adj_ < 0.001; Control vs JP, Mann-Whitney *U*-test, claudin-7: Z = 11.04, p_adj_ < 0.001; β-catenin: Z = 5.05, p_adj_ < 0.001). Unexpectedly, latrunculin A, which promotes filament depolymerization by sequestering G-actin, enhanced the membrane localization of both claudin-7 and β-catenin (Figure 3 A-C). In contrast, jasplakinolide, which inhibits F-actin disassembly, induced disintegration of the cortical network and the consequent delocalization of claudin-7 and β-catenin from the lateral membrane (26) (Figure 3D). These opposing outcomes demonstrate that the stability of tight and adherens junctions is not merely a passive consequence of F-actin integrity, but is instead differentially regulated by the specific nature of actin dynamics disruption. ROCK inhibition disrupted F-actin and resulted in the loss of claudin-7 from the lateral membrane (Figure 3 I-J) (Control vs Y-27623, Mann-Whitney *U*-test, claudin-7: Z = 3.35, p_adj_ < 0.01). However, the effect of ROCK inhibition on junctional integrity is strongly context-dependent, producing outcomes distinct from those of latrunculin A or jasplakinolide (26,27).

Given the critical role of actin in supporting junctional complexes, we hypothesized that microfilament destabilization is a principal cause of epithelial barrier dysfunction. In intestinal organoids, LatA treatment significantly increased 4 kDa FITC-dextran permeability (Figure 3 G-H), demonstrating that disruption of the F-actin cytoskeleton is sufficient to compromise barrier integrity (one-way ANOVA, F(2,33) = 7.22, p < 0.01, Tukey’s post-hoc test, Control vs LatA p_adj_ < 0.01). These results support our hypothesis that F-actin instability may be key to junctional dysfunction in *Muc2* KO mice.

### F-actin and junctional complexes are impaired in other chronic colitis models

We next sought to determine whether F-actin destabilization is a common pathogenic driver by examining two additional models of chronic colitis: DSS-induced and adoptive transfer colitis (ATC) (28,29). Fluorescence quantification confirmed that both models recapitulate the cytological defects observed in *Muc2* KO mice, namely disruption of actin cytoskeleton and altered claudin-7 localization (DSS vs Control, Mann-Whitney *U*-test, apical F-actin: Z = -15.03, p_adj_ < 0.001; lateral F-actin: Z = -11.38, p_adj_ < 0.001; claudin-7: Z = -9.18, p_adj_ < 0.001; ATC vs Control, Mann-Whitney *U*-test, apical F-actin: Z = -11.57, p_adj_ < 0.001; lateral F-actin: Z = -15.65, p_adj_ < 0.001; claudin-7: Z = -11.45, p_adj_ < 0.001). Additionally, altered β-catenin localization was confirmed in the DSS-colitis model (Figure 4) (DSS vs Control, Mann-Whitney *U*-test, β-catenin: Z = -5.07, p_adj_ < 0.001; ATC vs Control, Mann-Whitney *U*-test, β-catenin: Z = -0.62, p_adj_ > 0.05). This consistent phenotype across genetically, chemically, and immunologically driven models strongly suggests that microfilament stability fundamentally regulates barrier function in chronic colitis. One of the questions that remain open is the upstream mechanism responsible for the actin cytoskeleton remodeling.

**Figure 4.**
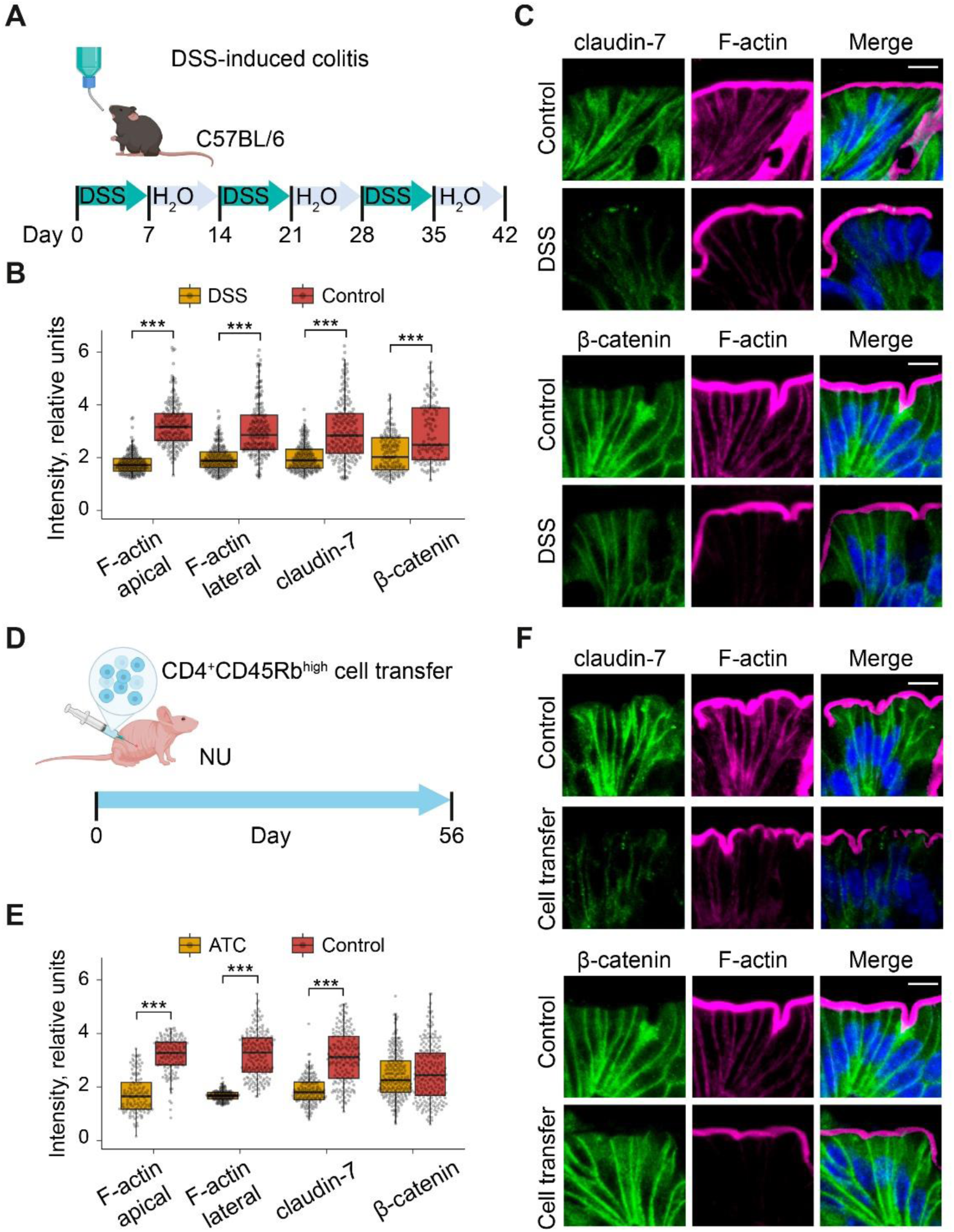
Actin dynamic is impaired in other chronic colitis models. (A) Scheme of chronic DSS-induced colitis development. (B) Fluorescence intensity quantification of F-actin, β-catenin and claudin-7 along the cell membrane in the descending colon of mice with DSS-induced chronic colitis (*** *p* < 0.001, n = 3, Mann-Whitney *U*-test). (C) Immunostaining of β-catenin and claudin-7 in the descending colon of mice with chronic colitis induced by DSS. Bar ‒ 10 µm. (D) Scheme of CD4^+^CD45Rb^High^-cells ATC development. (E) Fluorescence intensity quantification of F-actin, β-catenin and claudin-7 along the cell membrane in the descending colon of mice with CD4^+^CD45Rb^High^-cells ATC (*** *p* < 0.001, n = 3, Mann-Whitney *U*-test). (F) Immunostaining of β-catenin and claudin-7 in the descending colon of mice with ATC. Bar ‒ 10 µm.

### Lipid metabolism is implicated in barrier dysfunction in chronic colitis models

Given that metabolic reprogramming is common in IBD patients, we hypothesized that similar alterations in *Muc2* KO mice drive microfilament instability and junctional impairment. We conducted a comprehensive metabolomic profiling of isolated colonic crypts using liquid chromatography with tandem mass-spectrometry. Principal component analysis of metabolomes demonstrated significant differences between *Muc2* KO and wild-type littermates (F = 4.04, R² = 0.336, p = 0.009, 999 permutations), while intergroup comparisons revealed hundreds of differentially-regulated metabolites indicating a significant metabolic shift upon colitis (Figure 5 A, B). Metabolomic analysis revealed significant dysregulation of purine and pyrimidine metabolism, fatty acid biosynthesis, phospholipid, and sphingolipid metabolism (Figure 5 C, D). Sphingolipids were markedly elevated, with a notable accumulation of ceramides and glycosphingolipids (Figure 5D). Each lipid group represents a wide range of compounds with varying fatty acid length, saturation, and modifications. In particular, significant changes were seen across a wide spectrum of lipid species defined by their head groups and varying fatty acid chains. We detected significant alterations in 169 ceramides, 117 phosphatidylcholines, 102 phosphatidylglycerols, 43 phosphatidylserines, 25 phosphatidylinositols, 27 sphingomyelins, and 17 phosphatidic acids. Notably, the most significantly upregulated ceramides were enriched in long-chain (C16-C19) and very-long-chain (C20-C24) fatty acid classes, with a subset of ultra-long-chain ceramides (>C24) (Figure 5E).

**Figure 5.**
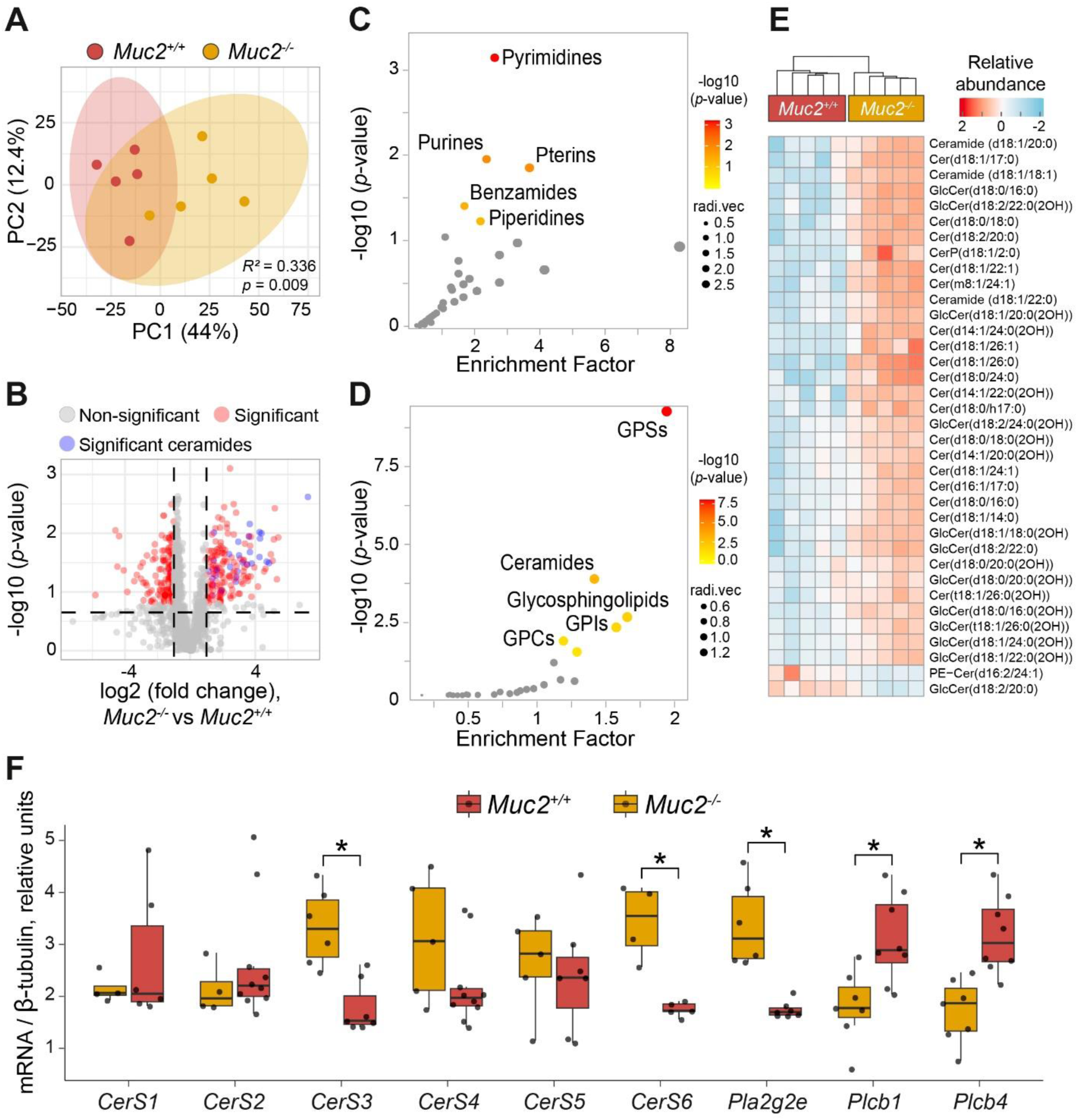
Phospholipid metabolism is impaired in the *Muc2* KO chronic colitis model. (A) Principal component analysis of metabolomic profiles in the colonic crypts of *Muc2* KO and littermate controls. (B) Volcano-plot showing significant increase of ceramides in *Muc2* KO crypts. (C) Small molecule enrichment analysis of metabolomic profiles obtained from *Muc2* KO crypts using HMDB database. (D) Lipid enrichment analysis of metabolomic profiles obtained from *Muc2* KO crypts using Lipid Maps database. GPCs ‒ glycophosphocholines, GPIs ‒ glycophosphoinositides, GPSs ‒ glycophosphoserines. (E) Heatmap showing abundance of different ceramides in *Muc2* KO crypts. (F) Expression analyses of genes associated with lipid metabolism in colonic tissues; mRNA levels were normalized on *Tubb5* (β-tubulin) using the deltaCt method. Resulting data were mean-centered and compared using Mann-Whitney *U*-test (* *p* < 0.05, n = 7-8).

Guided by our transcriptome data (not shown), we measured the expression of specific genes implicated in the regulation of F-actin dynamics and lipid metabolism. The expression of these candidates was quantified by RT-qPCR in an independent animal cohort to confirm their dysregulation. Genes encoding phospholipases A and C significantly change expression (*Muc2^-/-^*vs *Muc2^+/+^*, Mann-Whitney *U*-test, *Pla2g2e*: Z = 3, p_adj_ < 0.05; *Plcb1*: Z = -2.77, p_adj_ < 0.05; *Plcb4*: Z = -3, p_adj_ < 0.05) (Figure 5F). Interestingly, ceramide synthases 3 and 6 are significantly upregulated upon colitis in *Muc2*-deficient mice (*Muc2^-/-^* vs *Muc2^+/+^*, Mann-Whitney *U*-test, *CerS3*: Z = 2.86, p_adj_ = < 0.05; *CerS6*: Z = 2.45, p_adj_ < 0.05) (Figure 5F). The integration of gene expression and metabolomic data suggests that lipid accumulation might be associated with the disrupted cytoskeletal dynamics. Given the published data on the association of ceramides with poor IBD prognosis (30) and their upregulation in *Muc2* KO, we focused on ceramides as the leading candidate metabolites potentially mediating F-actin integrity.

### Inhibition of ceramide biosynthesis rescues intestinal barrier function

To functionally test the hypothesis that ceramide accumulation disrupts epithelial barrier, we inhibited ceramide synthesis using fumonisin B1 (FB1). Fumonisin B1, an inhibitor of ceramide synthases and a known toxin, was used in low doses via rectal administration. This treatment partially restored F-actin organization in *Muc2* KO mice (FB1 vs *Muc2^-/-^*, Mann-Whitney *U*-test, apical F-actin: Z = 6.24, p_adj_ < 0.001; lateral F-actin: Z = 4.37, p_adj_ < 0.001) (Figure 6 A, B). Inhibition of ceramide biosynthesis also affected the distribution of junctional proteins, promoting the lateral localization of β-catenin while displacing claudin-7 (FB1 vs *Muc2^-/-^*, Mann-Whitney *U*-test, β-catenin: Z = 2.31, p_adj_ < 0.05; claudin-7: Z = -4.9, p_adj_ < 0.001) (Figure 6 A, B). *In vivo* functional permeability test showed that rectal administration of fumonisin B1 reduced serum FITC-dextran 4 kDa levels in mucin-deficient animals, indicating at least partial restoration of the intestinal barrier function (FB1 vs *Muc2^-/-^*, Mann-Whitney *U*-test, Z = -2.43, p < 0.05) (Figure 6C). In DSS-treated mice, fumonisin B1 exposure also resulted in a significantly increased F-actin fluorescence in colon sections (FB1 vs DSS, Mann-Whitney *U*-test, apical F-actin: Z = 7.79, p_adj_ < 0.001; lateral F-actin: Z = 6.87, p_adj_ < 0.001) (Figure 6 D, E), accompanied by β-catenin mislocalization, but not claudin-7 (FB1 vs DSS, Mann-Whitney *U*-test, β-catenin: Z = 2.43, p_adj_ < 0.05; claudin-7: Z = -0.36, p_adj_ > 0.05) (Figure 6E). These data indicate that the mechanism of chronic inflammation also involves sphingolipid metabolism, which interferes with F-actin dynamics and the stability of intercellular junctions.

**Figure 6.**
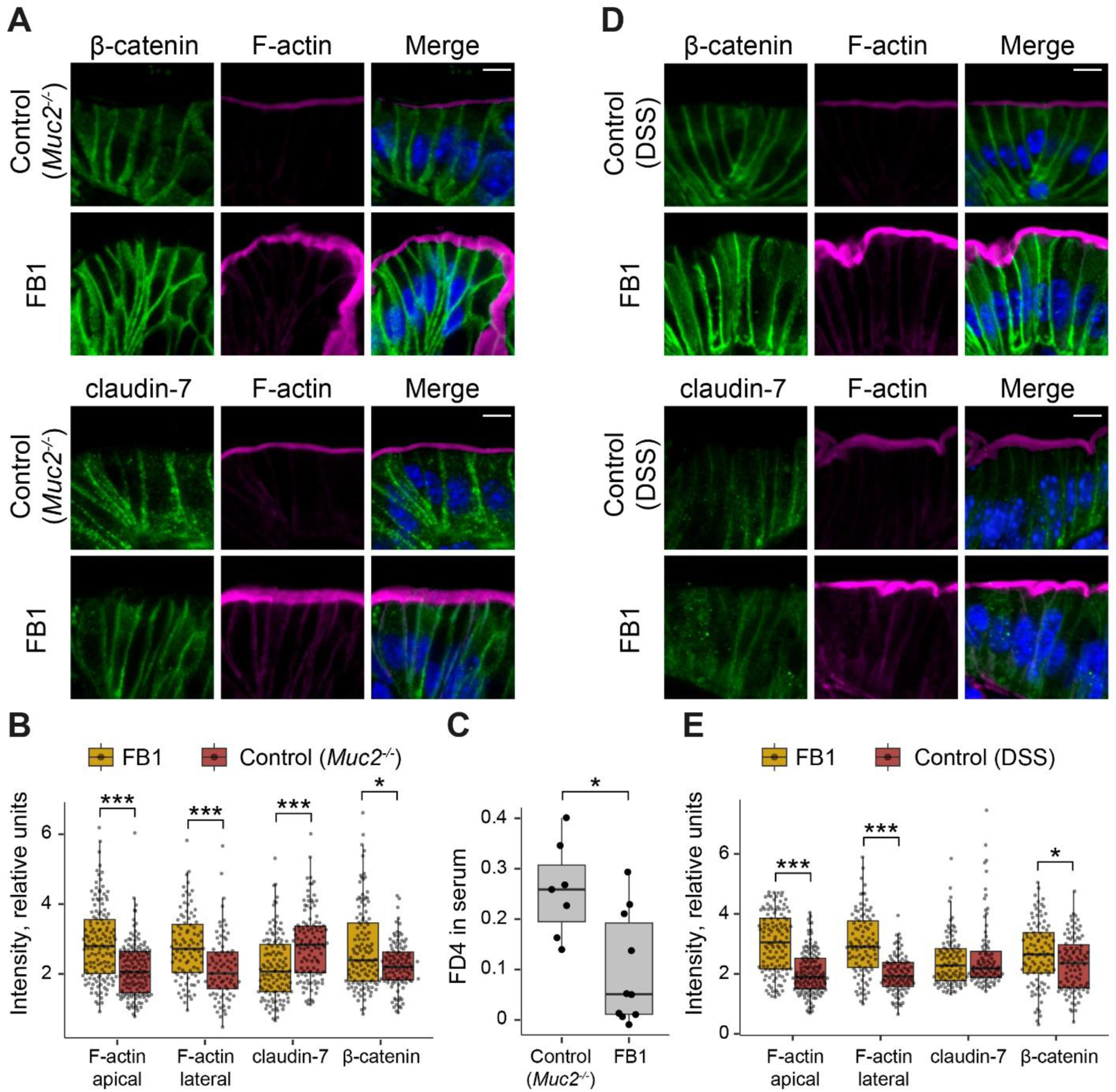
Inhibition of ceramide biosynthesis rescues intestinal permeability in mouse models of chronic colitis. (A) Claudin-7 and β-catenin immunostaining in the descending colon of *Muc2* KO mice after rectal administration of fumonisin B1 (FB1) for 12 hours. Bar ‒ 10 µm. (B) Fluorescence intensity quantification of F-actin, claudin-7 and β-catenin along the cell membrane of *Muc2* KO gut treated by fumonisin B1 (Mann-Whitney *U*-test, *** *p* < 0.001, n = 3). (C) *In vivo* permeability assay in *Muc2* KO mice upon fumonisin B1 treatment (Mann-Whitney *U*-test, * *p* < 0.05, n = 7-10). (D) Claudin-7 and β-catenin immunostaining in the descending colon of mice with DSS-induced colitis after rectal administration of fumonisin B1 for 12 hours. Bar ‒ 10 µm. (E) Fluorescence intensity quantification of F-actin, claudin-7 and β-catenin along the cell membrane of gut with DSS-induced colitis treated by fumonisin B1 (Mann-Whitney *U*-test, *** *p* < 0.001, n = 3).

### F-actin dynamics and TJ and AJ organization in patients with UC

To validate our findings in human tissues, we performed immunohistochemical analysis and transmission electron microscopy (TEM) on biopsies obtained from patients with ulcerative colitis (UC) and healthy controls. Fluorescence imaging of epithelial F-actin in colonic sections revealed a marked reduction of apical and lateral microfilaments, indicating pronounced cytoskeletal reorganization in UC patients (CTR_Colon vs UC_Colon, Mann-Whitney *U*-test, apical F-actin: Z = -26.82, p_adj_ < 0.001; lateral F-actin: Z = -29.85, p_adj_ < 0.001; UC_Colon vs UC_Ileum, apical F-actin: Z = 26.62, p_adj_ < 0.001; lateral F-actin: Z = 13.86, p_adj_ < 0.001) (Figure 7 A, B). TEM examination showed disrupted TJs and AJs in colonic samples from UC patients compared with ileal junctions from the same individuals and colonic tissue from non-IBD controls (Figure 7C). Morphometric assessment confirmed significant shortening of both TJs and AJs accompanied by an increased paracellular space within junctional regions (CTR_Colon vs UC_Colon, Mann-Whitney *U*-test, TJ length: Z = 1.58, p_adj_ = 0.11; TJ width: Z = 6.03, p_adj_ < 0.001, AJ length: Z = 7.46, p_adj_ < 0.001; AJ width: Z = 2.45, p_adj_ < 0.05 ; UC_Colon vs UC_Ileum, TJ length: Z = -3.91, p_adj_ < 0.001; TJ width: Z = -5.55, p_adj_ < 0.001, AJ length: Z = -3.1, p_adj_ < 0.01; AJ width: Z = -6.88, p_adj_ < 0.001) (Figure 7D). Together, these data demonstrate substantial impairment of the F-actin and junctional complexes in the colonic epithelium of patients with UC.

**Figure 7.**
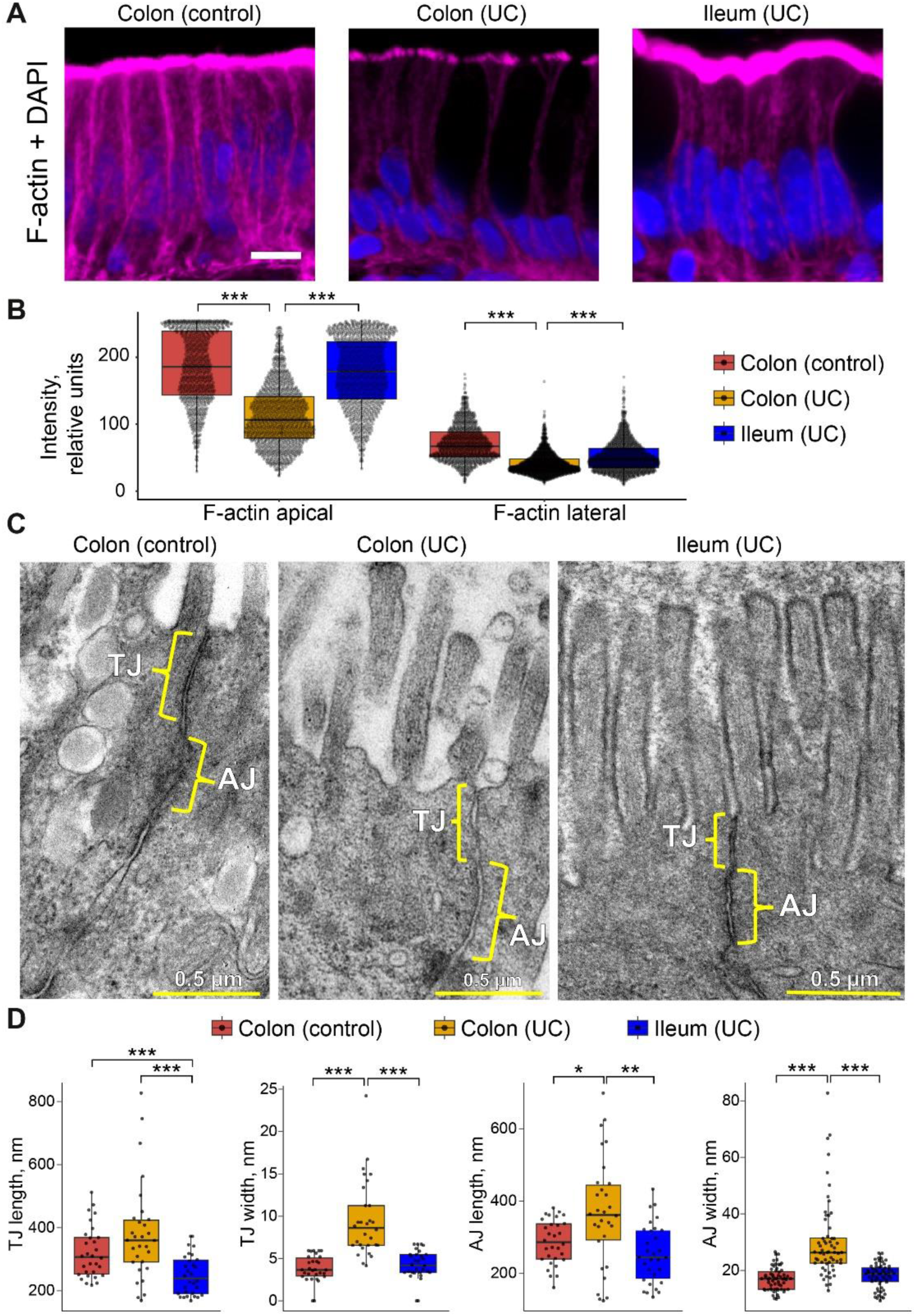
Intestinal barrier in patients with UC. (A) F-actin staining in IBD colonic and ileal biopsies and healthy colonic tissue. Bar ‒ 10 µm. (B) Fluorescence intensity quantification of F-actin along the cell membrane in IBD patients (Mann-Whitney *U*-test, *** *p* < 0.001, n = 3 IBD patients, n = 2 healthy controls). (C) TEM images of apical junctional complexes between enterocytes. TJ ‒ tight junction, AJ ‒ adherens junction. (D) Morphometric quantification of length and width of TJs and AJs (Mann-Whitney *U*-test, *** *p* < 0.001, n (UC patients) = 3, n (non-IBD controls) = 2).

## Discussion

### Actin dynamics and its interplay with junctional complexes

Multiple studies investigating the expression and localization of TJ and AJ proteins have focused on acute inflammation in both animal models and patients. Generally, these reports describe a downregulation of JAM-A, ZO-1, claudin-7, E-cadherin, and β-catenin (17,18,31–38). There are also report that demonstrate consistent or increased levels of TJ and AJ proteins in inflammation highlighting the complex nature of IBD (34,37,39,40). Here, we describe the delocalization of claudin-7, β-catenin, and E-cadherin from the lateral plasma membrane without changes of the total protein levels. These changes are accompanied by substantial F-actin downregulation across three different *in vivo* colitis models and in UC patients. Mechanistically, this delocalization is likely driven by endocytosis (41). For example, pro-inflammatory cytokines such as TNFα and IFNγ are known to increase epithelial permeability (38,42). Studies indicate this effect is mediated by the internalization of junctional proteins, as blockade of endocytosis preserves barrier function (38,41,42).

While the physical association between F-actin and junctional complexes (AJ and TJ) is well-established in healthy epithelial tissues, the fate of this interaction during IBD pathogenesis remains unclear. Substantial evidence indicates that inflammatory mediators disrupt actin cytoskeleton in model intestinal epithelial monolayers. However, the data regarding F-actin reorganization during *in vivo* inflammation are inconsistent and controversial (19,43,44). In this study, we demonstrate that the actin cytoskeletal network constitutes the majority of the claudin-3 interactome, demonstrating a profound physical interaction. Importantly, this interaction is lost in chronic colitis. Consistently, we show that targeting the actin monomer-polymer balance using latrunculin A or jasplakinolide not only disassembles F-actin in enterocytes but also concomitantly impairs the localization and function of TJs and AJs. These results agree with our *in vivo* models and UC patients’ data and with previous studies reporting that actin dynamics is linked to barrier function in cell cultures (45–47). Thus, investigating actin architecture in the extended cohort of UC and in CD patients using precision microscopy is a promising strategy. Alternatively, testing F-actin modifying drugs as potential barrier treatment represents another promising future direction.

### Ceramides and barrier function

Intestinal inflammation is associated with elevated ceramide levels (48–51). Ceramide levels increase by 71% in the DSS-induced chronic colitis model, and up to 160% in ATC models (49). As we report here, *Muc2* KO model is also characterized by upregulated intestinal sphingolipids, including ceramides. This result is consistent with *CerS3* and *CerS6* upregulation in the colon of *Muc2* KO mice. Notably, CerS3 is specific to the skin (52,53) and is specialized on producing very-long-chain ceramides (53), which aligns with our metabolomic profiling in *Muc2* KO mice and with data from IBD patients (54,55). Overexpression of CerS6 producing C14-C16 ceramides facilitates apoptosis in colonic adenocarcinoma cells (53). Elevation of C16 ceramides due to *CerS2* knockout exacerbates DSS-induced colitis (56).

We suggest that ceramide accumulation in the colonic epithelium disrupts actin cytoskeleton, leading to mislocalization of TJ/AJ proteins and a consequent increase in paracellular permeability. As secondary messengers, ceramides activate their downstream effectors ‒ protein phosphatase and atypical PKCζ ‒ which in turn modulate MAPK signaling to promote inflammatory and apoptotic responses (57). Ceramide-associated PKCζ regulates actin assembly and cell adhesion providing the link to cytoskeletal dynamics (58–60). Our data support the work of Bock et al., who found that SMase-induced ceramide accumulation in membrane fractions containing occludin and claudin-4 increases monolayer permeability, positioning ceramide as a primary inducer of barrier dysfunction (61). Furthermore, Kim et al. showed that *CerS2* knockout, which alters ceramide profiles, affects myosin light chain phosphorylation, thereby destabilizing actin filaments and increasing permeability (56).

Pharmacological inhibition of ceramide biosynthesis with fumonisin B1 partially restored barrier function in *Muc2* KO mice, likely by reducing pathogenic ceramide species. Ceramides can alter plasma membrane organization by accumulating in lipid rafts and displacing cholesterol which is associated with TJ dislocation and increased permeability (61–63). The interplay between lipid rafts and the actin cytoskeleton is bidirectional: raft composition can trigger Rho-dependent actin remodeling (64,65), while raft stability itself depends on F-actin (66). Actin depolymerization destabilizes rafts, redistributes raft-associated proteins, and enhances endocytosis (67). Our findings suggest that excessive ceramide accumulation impacts cytoskeletal reorganization and subsequent junctional defects, possibly via lipid raft remodeling. This potential mechanism is consistent with the study showing raft disruption in inflammatory conditions following cytokine exposure (68).

Another possible mechanism involves sphingosine metabolism. Sphingosine promotes the membrane association of β-catenin in cancer models and cell cultures (69). Ceramide synthases utilize sphingosine as a substrate possibly contributing to the delocalization of β-catenin protein (70). However, despite significant barrier recovery in *Muc2* KO mice, no changes in claudin-7 localization were observed. The precise role of claudin-7 remains unclear, as it has been reported to function as both a sealing and pore-forming protein (71,72). Given the context-dependent functional variability of claudin-7 (73), our result may indicate a possible role for other proteins of the claudin family.

It worth noting that fumonisin B1 is a potent mycotoxin unsuitable for clinical use (74) and its effects strongly depend on the route of administration. While our localized rectal delivery achieved a targeted effect, other studies reported increased epithelial permeability following oral fumonisin B1 treatment, likely due to prolonged exposure throughout the gastrointestinal tract (75–77). Nevertheless, our data provide proof-of-concept that inhibiting ceramide biosynthesis is a viable strategy for restoring the epithelial barrier in IBD. Given the toxicity of fumonisin B1, future efforts should focus on developing novel, low-toxicity precision inhibitors that can safely target this pathway.

## Conclusion

By analyzing three mouse models of chronic colitis with distinct etiology alongside colonic tissues from UC patients, we show that disruption of the F-actin cytoskeleton is the pivotal event in epithelial barrier dysfunction. The impaired F-actin dynamics is prerequisite to junctional complexes delocalization from the lateral plasma membrane leading to the TJ and AJ functional defects. One of the mechanisms behind F-actin network disruption involves lipid metabolism, with ceramides emerging as key molecules influencing cytoskeletal dynamics and intestinal permeability in chronic colitis. Thus, targeting the ceramide–actin interplay presents a novel therapeutic strategy for restoring epithelial integrity in IBD.

## Acknowledgements

We thank Dr. A.E. Ogienko and Dr. L.V. Boldyreva for technical support with microscopic analyses and sample management, T. N. Belovezhets for technical support with cell sorting, Dr. E. A. Stekolschikova and N. A. Anikanov for performing the LC-MS/MS analysis. We thank the Microscopic Center of the Siberian Branch of the Russian Academy of Sciences (FWNR-2022-0015), the "Recombinant antibody design" core facility of the IMCB SB RAS, and the Skoltech Advanced Mass Spectrometry Core Facility. We acknowledge the use of generative AI during the final editing phase to improve the manuscript’s readability and language.

## Author Contributions

Conceptualization, E.K..; methodology, S.M., J.P., K.A., J.K., K.M., and E.K.; formal analysis, S.M., J.P., K.A., L.S., and E.K.; investigation, S.M., J.P., K.A., J.K, E.N., K.M., M.O., and E.K.; resources, J.K. and E.K.; data curation, K.A. and E.K.; writing – original draft preparation, S.S., and E.K.; writing – review and editing, S.M., J.P., K.M., and E.K.; visualization, S.M., J.P., and E.K.; supervision, E.K.; project administration, E.K.; funding acquisition, E.K. All authors have read and agreed to the published version of the manuscript.

## Data availability

The metabolomic and proteomic data supporting the findings of this study are available for the reviewers from the corresponding author upon request, and will be made publicly available at the time of publication. The study was conducted based on a detailed research plan that was peer-reviewed and funded as a grant proposal. The funded grant proposal is available from the corresponding author upon reasonable request.

## Materials & Methods

### Experimental model and subject details

#### Human subjects

This study included 3 adult patients with a confirmed diagnosis of ulcerative colitis (UC) and 2 healthy controls. Patient 1 was a 41-year-old female of European descent. The disease was classified as left-sided colitis (E2 according to the Montreal classification) with a chronic relapsing course. At the time of the study, the disease activity index (DAI) score was 6. The disease duration was 11 years. There was no family history of UC. Her baseline therapy consisted of Guselkumab for 4 years and azathioprine for 7 years. Her medical history included more than three courses of corticosteroid treatment, with the last course administered 6 years prior to the study.

Patient 2 was a 36-year-old male of European descent. The disease was classified as extensive colitis (pancolitis, E3 according to the Montreal classification) with a chronic relapsing course. At the time of the study, the patient was in clinical remission with the DAI score of 7. Extraintestinal manifestations included primary sclerosing cholangitis, aphthous stomatitis. The disease duration was 7 years. There was no family history of UC. His baseline therapy consisted of Guselkumab for 4 years and azathioprine for 5 years. His medical history included corticosteroid treatment, with the last course administered 6 months prior to the study.

Patient 3 was a 35-year-old male of European descent. The disease was classified as extensive colitis (pancolitis, E3 according to the Montreal classification) with a chronic continuous course. The patient had steroid-dependent disease. At the time of the study the DAI score was 8. Extraintestinal manifestations included primary sclerosing cholangitis at the cirrhotic stage, Sweet’s syndrome, and recurrent aphthous stomatitis. The disease duration was 7 years. There was no family history of ulcerative colitis. Baseline therapy consisted of biologic treatment with Guselkumab for 4 years and azathioprine for 6 years. The patient had a history of more than three courses of corticosteroid therapy; the study was conducted during ongoing corticosteroid treatment (prednisolone 40 mg).

Control subjects were 2 females (32 and 66-year-old) of European descent receiving screening colonoscopies without IBD diagnosis. Both participants reported no family history of IBD.

Biopsy samples were collected for analysis during a standard endoscopic procedure. From each patient, one sample was taken from the rectum and one from the ileum specifically for analysis by TEM and immunohistochemistry. Additional samples were collected for other analyses not reported here.

All procedures were reviewed and approved by the Ethics Committee of Novosibirsk State Medical University (Novosibirsk, Russia), protocol #164, 24 February 2025. All research was performed in accordance with the Declaration of Helsinki. Written informed consent was obtained from all participants involved in the study.

#### Mouse models

All procedures were conducted under Russian legislation according to the standards of Good Laboratory Practice (directive # 267 from 19 June 2003 of the Ministry of Health of the Russian Federation), institutional Ethical committee guidelines and the European Convention for the protection of vertebrate animals. All procedures were approved by the Ethical committee at IMCB (protocol #002 of 23.04.2025, #003 of 24.06.2025 and protocol #003 of 16.09.2024). All animals were quarterly tested for the specific pathogen free (SPF) status according to Federation of European laboratory animal science association’s (FELASA) recommendations.

The study was conducted using C57BL/6JNskrc (C57BL/6J sub-colony, referred as C57BL/6), BALB/cNskrc (BALB/cJ sub-colony, referred as BALB/c), NU (NU/J sub-colony) that were bred in the Institute of Cytology and Genetics SB RAS, and *Muc2^−/−^* mice that were bred in the Institute of Molecular and Cellular Biology SB RAS. *Muc2^−/−^* mice were bred as a subcolony of previously generated *Muc2^tm1Avel^/Muc2^tm1Ave^*^l^ mice on C57BL/6 genetic background. Mutant mice and their wild-type littermates (*Muc2^+/+^* mice) were obtained by crossing *Muc2^+/−^* females to *Muc2^+/−^*males. *Muc2^+/+^* mice were used as a control group in all experiments involving *Muc2^−/−^* mice to diminish the effect of microbiota. All experiments were performed in the Institute of Molecular and Cellular Biology SB RAS.

Adult mice were housed separately in open cages (with a dimension of 318 × 202 × 135 mm, #CP-3, 3 W, Russia) with birch sawdust as litter and paper cups as shelter. The housing conditions were as follows: 12 h/12 h light/dark photoperiod; and food (BioPro, Novosibirsk, Russia) and water were provided *ad libitum*. The 5-week-old offspring were weaned and placed in open cages in same-gender groups and housed under the same conditions. All experiments were carried out on mice between 8 and 12 weeks of age. Animals from the same cage were equally distributed among groups to avoid cage affect. Exclusion criteria from the experiment were weight loss of more than 20%; organic disorders of the central nervous system, alopecia, abscesses, injuries. Animals were euthanized via cervical dislocation. Descending colon samples were taken for immunofluorescence staining, RT-qPCR, metabolomic and transcriptomic analysis and crypts isolation.

#### Chronic DSS colitis

To induce chronic intestinal inflammation 12-week-old C57BL/6 mice (n = 6 per group) were treated with dextran sodium sulfate (DSS, #1138GR250, NeoFroxx, Germany) in drinking water. Mice received three cycles of DSS/Water (7 days 2% DSS followed by 7 days of water), control mice obtained drinking water. Mice were weighted three times a week for weight loss control. Chronic DSS treatment induced loose bloody stool, ruffled and unkempt fur. After the treatment (42 days in total) mice were taken for subsequent experiments.

#### Adoptive transfer colitis model

Spleens obtained from donor BALB/c mice were homogenized in cold (4℃) PBS completed with 4% FBS filtered through 70 um strainers. Red blood cells were lysed with cold (4℃) ammonium chloride hypotonic buffer (0.8% NH^4^Cl, 0.084% NaHCO_3_, 0.037% EDTA) for 10 min at room temperature. Then cells were centrifuged at 400×*g*, 4℃ for 5 min and washed with 4% FBS in PBS (4℃) 2 times. After the last wash cells were resuspended in a sorting buffer (0.5% BSA, 2mM EDTA in PBS) to concentration of 10^8^ cells/ml. CD4-positive cells were sorted by negative selection using magnetic beads-based MojoSort Mouse CD4 T-Cell Isolation Kit (#480005, BioLegend) according to manufacturer’s recommendations. Then isolated cells were stained with FITC anti-mouse CD45RB Antibody (#103305, BioLegend) and PE/Cyanine7 anti-mouse CD4 Antibody (#100421, BioLegend) for 45 min on ice, washed with 4% FBS in PBS (4℃). CD4^+^CD45Rb^High^ cells with brightest 24% were sorted using Sony SH800 Cell Sorter, centrifuged at 400×*g*, 4℃ for 10 min and resuspended in sterile PBS (4℃) for injection to recipient NU male mice (n = 3), 0.5 x 10^6^ cells in 250 µl per mouse as described previously (78). Control NU mice (n = 3) were injected with sterile PBS. Mice were weighed initially before T-cell transfer (or PBS injection) and then two times a week. They were observed for clinical signs of inflammation: appearance, diarrhea, blood in stool and rectal bleeding. Recipient mice developed intestinal inflammation with diarrhea in 6 weeks after T-cell transfer. When loss of body weight exceeded 20% after transfer, the host mice were sacrificed. All tissues were subjected to pathological observations and cytokine profile analysis at the endpoint.

### Method details

#### Immunohistochemical analysis

The descending colon from mice (n = 3 per group) and human colonic biopsies (n = 3 IBD patients, n = 2 healthy controls) were fixed overnight in 10% neutral buffered formalin, then kept in 15% sucrose for 12 h and in 30% sucrose for another 12 h. After that samples were embedded in O.C.T. Compound (Sakura Finetek, USA) and frozen in liquid nitrogen. 40 μm (mice tissues) and 20 μm (human tissues) sections were prepared using 550 HM Microm cryostat (ThermoFisher Scientific, USA). The slides were washed three times for 5 min in PBS + 0.3 % Triton X-100 (PBST). Sections were incubated in PBST + 0.5% bovine serum albumin (BSA) for 2 h. Primary antibodies (Table 1) were incubated overnight at 4°C in PBST + 0.5% BSA, then washed three times for 5 min each with PBST. Secondary antibodies (Table 1) were incubated for 2 h at room temperature in PBST + 0.5% BSA and washed three times for 5 min each with PBST. Lastly, sections were incubated with 1 µM TRITC-Phalloidin (#P1951, Sigma, USA) and 0.15 µg/ml DAPI (#TC229, HiMedia, India) on PBS for 1 h. The colonic sections were mounted in 1:1 PBS:glycerol on slides under cover glasses. All microscopic slides from one experiment (all comparison groups per experiment) were prepared simultaneously using the same antibody dilution.

**Table 1.**
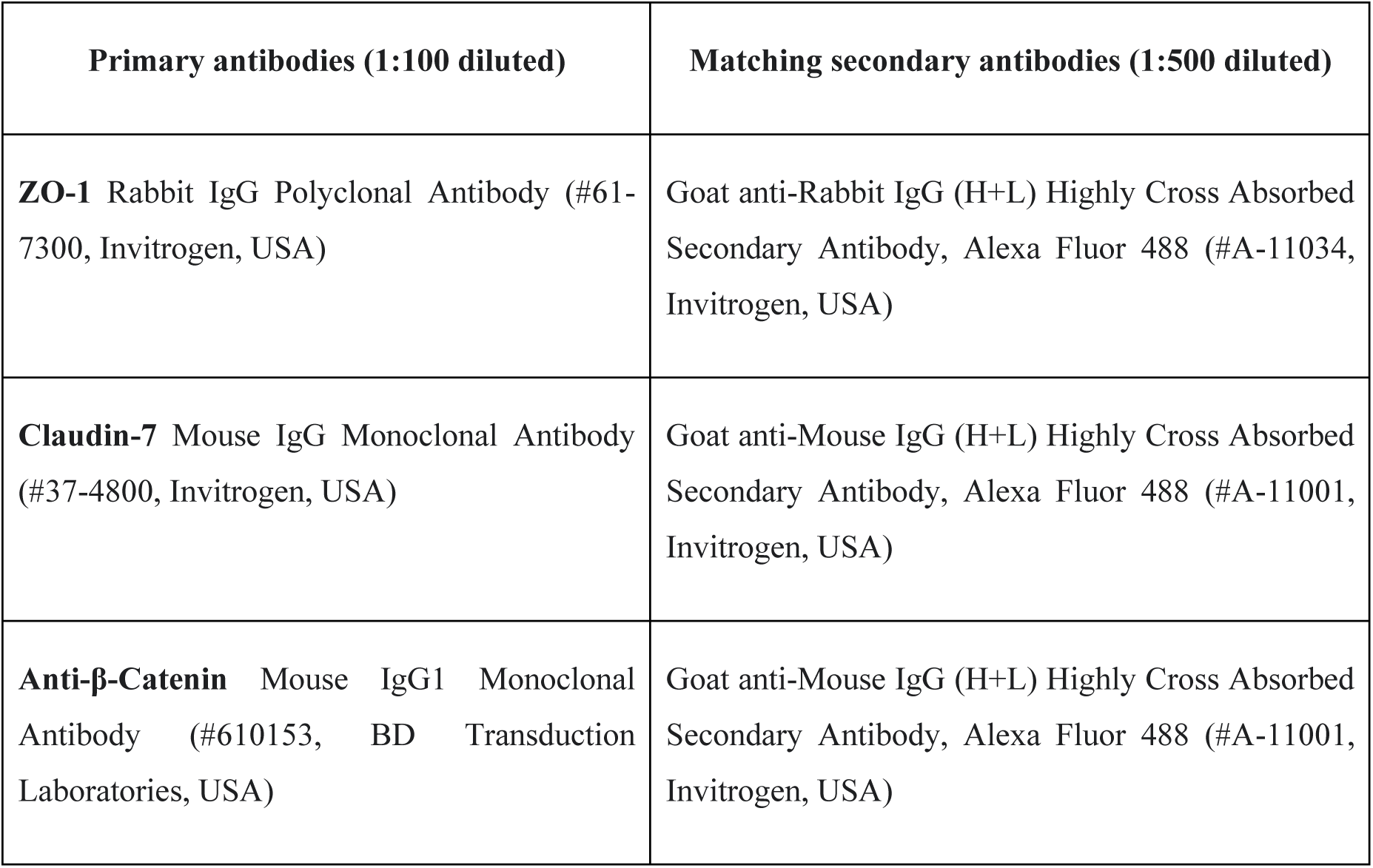
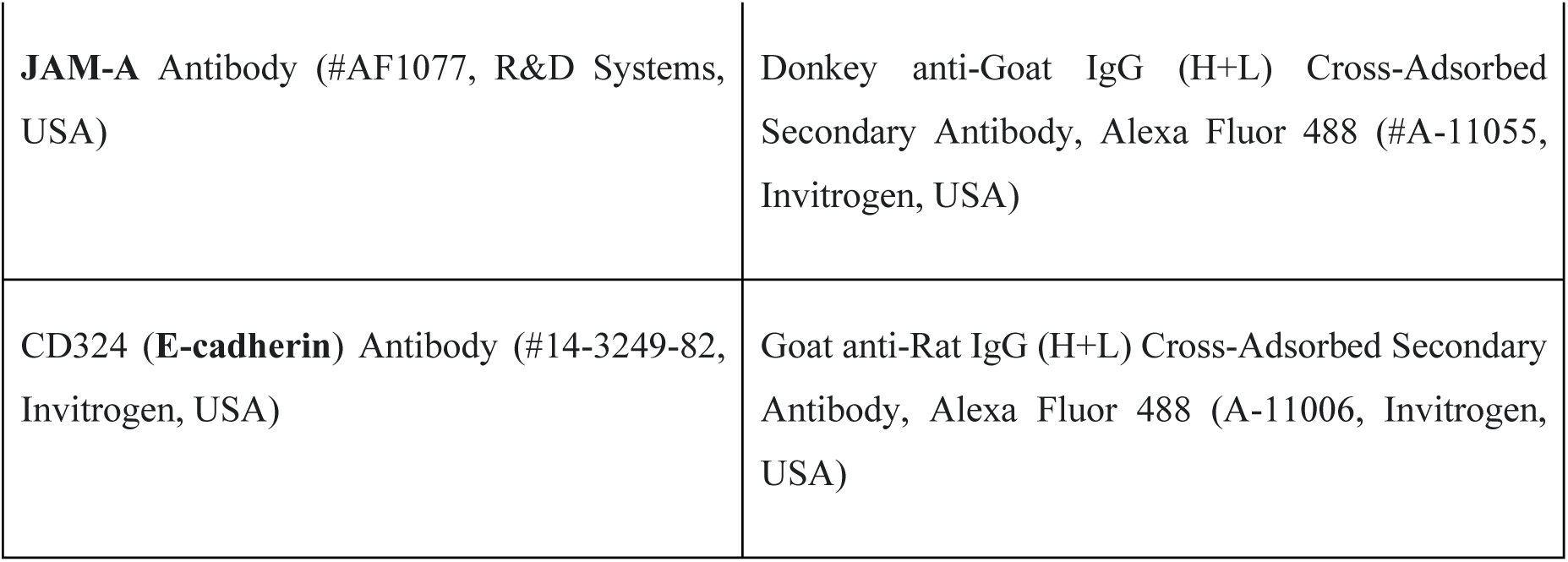
Antibodies used in the study for immunohistochemistry.

| Primary antibodies (1:100 diluted) | Matching secondary antibodies (1:500 diluted) |
| --- | --- |
| **ZO-1** Rabbit IgG Polyclonal Antibody (#61-7300, Invitrogen, USA) | Goat anti-Rabbit IgG (H+L) Highly Cross Absorbed Secondary Antibody, Alexa Fluor 488 (#A-11034, Invitrogen, USA) |
| **Claudin-7** Mouse IgG Monoclonal Antibody (#37-4800, Invitrogen, USA) | Goat anti-Mouse IgG (H+L) Highly Cross Absorbed Secondary Antibody, Alexa Fluor 488 (#A-11001, Invitrogen, USA) |
| **Anti-β-Catenin** Mouse IgG1 Monoclonal Antibody (#610153, BD Transduction Laboratories, USA) | Goat anti-Mouse IgG (H+L) Highly Cross Absorbed Secondary Antibody, Alexa Fluor 488 (#A-11001, Invitrogen, USA) |
| **JAM-A** Antibody (#AF1077, R&D Systems, USA) | Donkey anti-Goat IgG (H+L) Cross-Adsorbed Secondary Antibody, Alexa Fluor 488 (#A-11055, Invitrogen, USA) |
| **CD324 (E-cadherin)** Antibody (#14-3249-82, Invitrogen, USA) | Goat anti-Rat IgG (H+L) Cross-Adsorbed Secondary Antibody, Alexa Fluor 488 (A-11006, Invitrogen, USA) |

Images were obtained using a confocal microscope LSM 710 (Carl Zeiss, Germany) with oil immersion 63×/1.40 plan-apo objective and the ZEN 2012 software using the same settings for each experiment. Confocal microscopy was performed in the core facility of the Institute of Molecular and Cellular Biology SB RAS. 3D confocal slices of similar width (2.412 μm) reconstructed using maximal intensity method were saved as TIFF files for subsequent quantification and randomized by an independent researcher by assigning randomly generated filenames. Fluorescence intensity quantification was performed using ImageJ software using a freehand line tool by a single researcher not involved in data acquisition and randomization. Claudin-7 and β-catenin staining intensity was measured along the lateral cell membranes, F-actin staining was measured within the brush border as apical actin and along the lateral cell membranes as lateral actin. Lateral membranes were initially traced at the F-actin channel, then the corresponding area of TJ/AJ channel was measured. The background intensity was subtracted from every measurement. At least 30 independent measurements were made for each of the three biological replicates in the group. Image analysis was performed by a researcher blinded to the experimental group allocations.

#### Western blot analysis

For immunoblots, the descending colon samples (n = 6 per group) were homogenized in RIPA buffer (150 mM NaCl, 1% Nonidet P-40, 0.5% sodium deoxycholate, 0.1% SDS, 25 mM Tris (pH 7.4), 1 mM sodium metabisulphite, 1 mM dithiothreitol, 1 mM phenylmethylsulfonyl fluoride) using plastic pestles. The lysates were centrifuged at 12,000×*g* at 4℃ for 30 min, the supernatants were collected and used to quantify protein concentration as described by Bradford (79). Total protein extracts were boiled in SDS-PAGE sample buffer, and about 40 μg of total protein was loaded per lane of the 15% acrylamide gel. Mouse monoclonal anti-β-catenin (#610153, BD Biosciences, USA), mouse monoclonal anti-E-cadherin (#14-3249-82, Invitrogen, USA), mouse monoclonal anti-β-tubulin (#556321, BD Biosciences, USA) antibodies were used at 1:1000. Goat anti-mouse HRP (#G-21040 respectively, both ThermoFisher Scientific, USA, 1:3500) served as secondary antibodies. Images were captured using an Amersham Imager 600 System (GE Healthcare) and Novex ECL Chemiluminescent Substrate Reagent Kit (ThermoFisher Scientific, USA). Quantification of blots was performed using ImageJ software.

#### Intestinal organoids

Colon specimens were excised from mice, rinsed using 23 G catheter with cold (4℃) Dulbecco’s Phosphate Buffered Saline (DPBS) (#D5652, Sigma, USA) containing 1% antibiotic-antimycotic solution (ABM) (#A5955, Sigma, USA), then cut longitudinally and washed in cold DPBS+ABM 3 times. Colon specimens were cut in small pieces (to pass through 1 ml pipette tip) and digested in 2 mg/ml Collagenase I (#17018-029, Gibco, USA) solution in DMEM/F12 (#D9785, Sigma-Aldrich, USA) completed with 5% FBS (#SV30160.03, HyClone, USA) and Antibiotic-Antimycotic (#A5955, Sigma-Aldrich, USA) for 30 min at 37℃, 5% CO_2_. Crypts were released by pipetting with 1000 µl tip, filtering through a 70 µm strainer, then were centrifuged at 250×*g*, 4℃ for 5 min. The crypts were resuspended in Organoid-Star Matrigengel (#0827555, ABW, USA). The drops of 25 μl were seeded into 48-well culture plates. Plates were incubated for 5 min at 37℃ until the matrix polymerized. Culturing was performed in IntestiCult Organoid Growth Medium (#06000, Stemcell Technologies, Canada) with 1% ABM at 37℃ and 5% CO_2_. The medium was changed every three days. After 7-10 days of culture, organoids were passaged. For passaging, IntestiCult was removed, Matrigengel drops were resuspended in 0.25% trypsin-EDTA (#T4049, Sigma, USA) and incubated for 5 min at 37℃, 5% CO_2_. Organoids were washed with DMEM/F12 + 10% FBS + 1% ABM, centrifuged at 200×*g* for 5 min. The supernatant was removed, organoids were resuspended in Matrigengel and seeded as described earlier. Permeability assay was conducted after 3 days post passaging.

#### Permeability *in vitro* assay

Drops of Cultrex with 10-day organoids were destroyed with pipetting. 2 mg/ml FITC-Dextran 4 (#FD4-1G, Sigma, USA) and 1 mg/ml Hoechst 33342 (#G1127, ServiceBio, China) were added to each well for 30 min. For experimental wells also 1 µM jasplakinolide (#sc-202191, Santa Cruz Biotechnology, USA) or 2 µM latrunculin A (#100-0562, Stemcell Technologies, Canada) were added for 30 min. After incubation, organoids were transferred to the 8-well μ-slide (SPL Lifesciences, Korea) and observed under a confocal microscope LSM 710 (Zeiss) with the ZEN 2012 software. The μ-slide was placed on a temperature-controlled microscope stage (37℃) with CO_2_ supplied. The microscopy was conducted at the Joint Access Center for Microscopy of Biological Objects with the Siberian Branch of the Russian Academy of Sciences. Fluorescence intensity was quantified using ImageJ software. Permeability was evaluated as FD4 fluorescence intensity in the lumen of the organoid divided on the mean of three background regions around the organoid.

#### Intestinal permeability *in vivo*

Intestinal permeability was measured using 4 kDa FITC-Dextran (FD4) (#FD4-1G, Sigma, USA), n = 10 for control, n = 13 for experiment. A total of 100 μL FD4 (20 mg/mL in PBS) was administered by oral gavage using a steel feeding tube. After 4 h, 200 μL of blood was collected by decapitation. Blood was diluted with PBS containing 0.5% heparin and centrifuged at 250×*g* for 15 min at 4℃. A total 100 µL of the supernatant was applied to a 96-well plate, and FITC (485 nm / 535 nm) fluorescence was measured using TriStar LB 941 (Berthold Technologies, Germany). Baseline blood plasma fluorescence was determined in mice after oral gavage with water (n = 3 for each group) and subtracted from fluorescence obtained after FD4 gavage. FD4 concentrations were determined from standard curves generated by serial dilutions of FD4 in PBS.

#### Rectal administration of inhibitors

Mice (n = 3 per group) were anesthetized intraperitoneally (Domitor, 0.2 mg per1 kg body weight and Zoletil, 20 mg per1 kg body weight). Mice were rectally administered 15 µM latrunculin A (#100-0562, Stemcell Technologies, Canada), 3 µM Jasplakinolide (#sc-202191, Santa Cruz Biotechnology, USA), 20 µM Y-27623 (Stemcell Technologies, Canada) or 20 µM fumonisin B1 (#73682, Stemcell Technologies, Canada) dissolved in 33 µl of 0.9% NaCl. Control mice were given 33 µl of 0.9% NaCl. After that, mice were left lying on their back on the warm surface (37℃) with their tails raised to prevent enema leakage until awakened. Then animals were put back to their home cages. After 3 h (latrunculin A/jasplakinolide) or 12 h (Y-27623 or fumonisin B1) since administration mice were sacrificed and their descending colons were harvested for immunohistochemical analysis.

#### Metabolomic analysis

Colon specimens were excised from mice (n = 5 per group), rinsed using 23 G catheter with cold (4℃) Dulbecco’s Phosphate Buffered Saline (DPBS) (#D5652, Sigma, USA) containing 1% antibiotic-antimycotic solution (ABM) (#A5955, Sigma, USA), then cut longitudinally and washed in cold DPBS+ABM 3 times. Colon specimens were cut in small pieces (to pass through 1 ml pipette tip) and incubated in 3 mM EDTA (Sigma-Aldrich, USA) and 1 mM DTT (Sigma-Aldrich, USA) in PBS at 37℃ for 30 min. After vigorous shaking, the suspension was filtered through a 70 μm strainer. The crypt suspensions were precipitated (250×*g*, 5 min, 4℃), and the pellets were immediately frozen in liquid nitrogen. An additional aliquot (1/10 of the volume) was taken from each sample for subsequent total protein determination and normalization. Frozen samples were shipped on dry ice to the Skolkovo Institute of Science and Technology (Skoltech, Moscow, Russia) Advanced Mass Spectrometry Core Facility. Non-targeted metabolomic analysis was performed using high-performance liquid chromatography coupled with tandem mass spectrometry. Samples were cold homogenized, and metabolite extraction was performed with 80% methanol. The resulting extracts were dried, then redissolved in 20% acetonitrile, and separated by liquid chromatography.

Metabolites with annotations in HMDB (Human Metabolome Database) and Lipid Maps databases were filtered out of total metabolomics data obtained by LC-MS/MS, and analyzed using Metaboanalyst 5.0 online tool (80). The data were normalized by median, *log10*-transformed and Pareto scaling was applied. Then Fold Change analysis, t-test and PCA were performed. Additionally, KEGG Pathway Analysis on normalized by median and log10-transformed data was conducted. For significantly different annotated metabolites and differentially expressing genes (FDR > 0.05, Fold Change ≥ 2) Pathway Joint Analysis was performed.

#### Gene expression analysis

Distal colon tissue (about 5 mm) was collected and stored at -70℃ until the analysis. RNA was extracted using QIAzol Lysis Reagent (#79306, Qiagen, Germany) according to manufacturer’s recommendations. Purified RNA was treated with DNase I (#04716728001, Switzerland) and precipitated with ethanol. RNA concentrations were measured using Nanodrop 2000 (ThermoFisher Scientific, USA). Reverse transcription was performed using M-MuLV reverse transcriptase (SibEnzyme, Novosibirsk, Russia), a mix of random hexa-deoxyribonucleotide and Oligo-dT primers and 5 µg of RNA in reaction volume of 40 µl, after the reaction mixture was diluted up to 200 µl. Expression of the genes of interest was assessed by Real-time PCR using a BioMaster HS-qPCR SYBR Blue (2x) (BioLabMix, Novosibirsk, Russia), 250 nM specific primers (listed in Table 2), and 5 µl of cDNA in a CFX96 Touch Real-Time PCR Detection System (BioRad, Hercules, USA). Gene expression was normalized to *Tubb5* (tubulin, beta 5 class I) mRNA level as ΔCt = 2 ^ (Ct_Tubb5 mRNA_-Ct_gene of interest mRNA_).

**Table 2.**
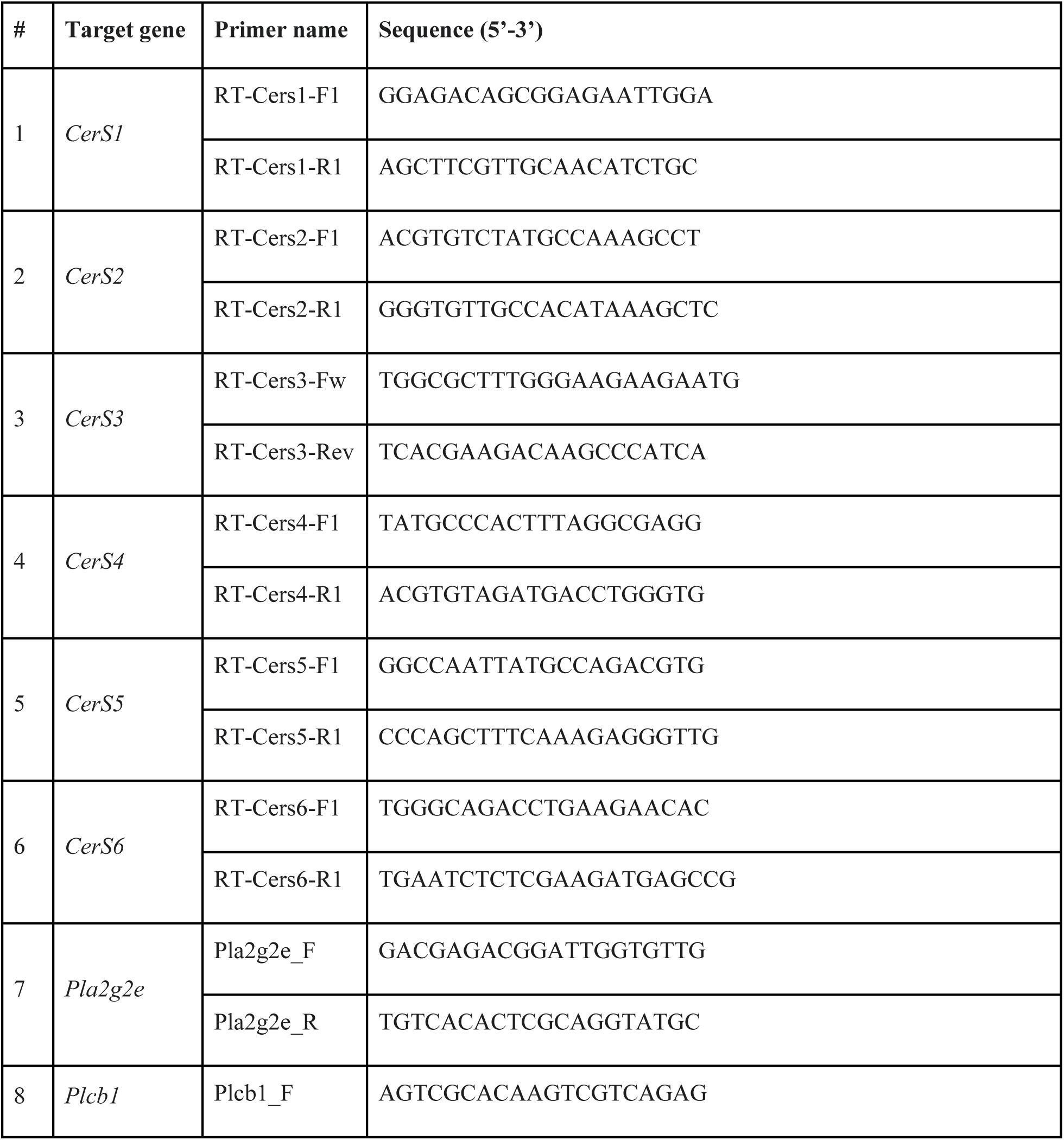
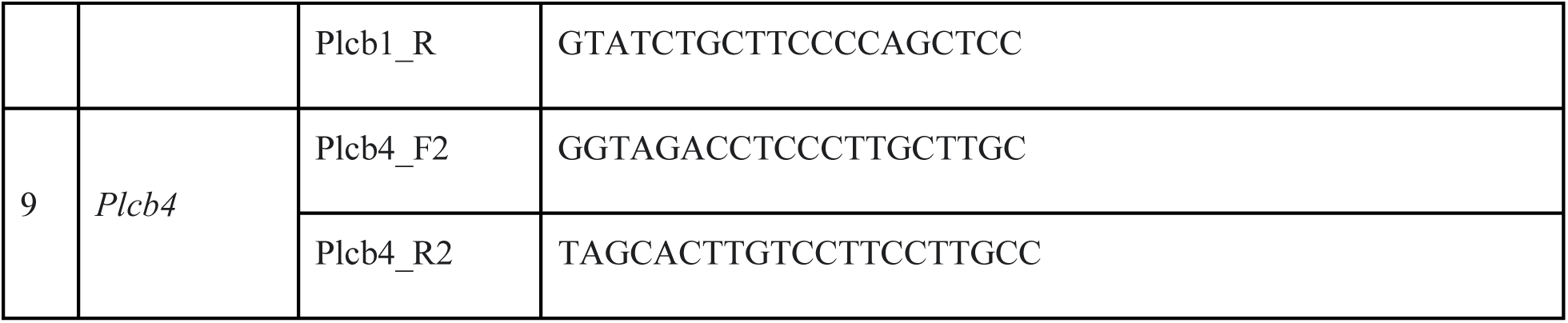
Primers used in the study.

#### Immunoprecipitation and proteomics

Colonic tissues were collected from *Muc2^+/+^* and *Muc2* KO mice (n = 6 per group) and immediately homogenized in ice-cold lysis buffer containing 25 mM HEPES (pH 8.0), 200 mM NaCl, 5 mM EDTA, 0.1% Triton X-100, supplemented with 1 mM sodium metabisulfite (NAMBS) and 1 mM phenylmethylsulfonyl fluoride (PMSF). Homogenization was performed using a glass homogenizer with a tight pestle. Lysates were centrifugated at 13,000×*g* for 30 min at 4℃, and the supernatants from each genotype were combined for immunoprecipitation. For each genotype, 2.5 mg of total protein extract was incubated with 50 µL of anti-claudin-3 antibody (#ab15102, Abcam, UK) and Protein A Sepharose beads (70-80 µL packed volume). Binding reactions were carried out in 25 mM HEPES (pH 8.0), 200 mM NaCl, 5 mM EDTA, 0.1% NP-40, 1 mM NAMBS, and 1 mM PMSF for 4 h at 4℃. Beads were washed three times with a wash buffer (25 mM HEPES, pH 8.0, 500 mM NaCl, 5 mM EDTA, 0.1% NP-40, 1 mM NAMBS, 1 mM PMSF). Protein complexes were eluted in citrate buffer (pH 3.0), and eluates were immediately neutralized. Protein precipitation was performed using the trichloroacetic acid / deoxycholate (TCA/DOC) method. Pellets were resuspended in 35 µL of Laemmli sample buffer and heat-denatured prior to proteomic analysis. A mock IP control was performed using non-functional mock antibodies (verified to show no reactivity with colonic lysates in Western blot). For this control, a 1:1 mixture of protein extracts from *Muc2* KO mice and their wilt-type littermates was used. Denatured samples were shipped to BGI (Hong Kong, China) for LC-MS/MS analysis and peptide annotation.

Intensity-Based Absolute Quantification (iBAQ) scores for proteins identified in the mock IP were subtracted from the corresponding values from the IPs from *Muc2* KO mice and the control animals. Next, protein lists were depleted for proteins annotated to heat shock proteins, immunoglobulins, keratins, ribosomal proteins and elongation factors as these proteins are common nonspecific contaminants in immunoprecipitation experiments and are unlikely to be related to tight junctions. The remaining protein lists were subjected to functional annotation (gene ontology, GO) using clusterProfiler (81) and org.Mm.eg.db (82) R packages based on “Biological process” ontology.

#### Transmission Electron Microscopy

Human biopsy samples (both ileum and colon, n = 3 IBD patients, n = 2 healthy controls) were placed in a 2.5% glutaraldehyde solution in a 0.1 M sodium cacodylate buffer (pH 7.4) for 24 h at room temperature, washed and post-fixed in 1% osmium tetroxide with 0.8% potassium ferrocyanide for 1 h. Fixed samples were contrasted with 1% uranyl acetate in mQ water, dehydrated and embedded in epoxy resin (Epon 812). Semi-thin cross-sections were prepared, stained with 1% methylene blue and analyzed with an Axioscope-4 microscope (Carl Zeiss). Ultrathin (70 nm) sections for TEM were obtained with a diamond knife (Diatome, Nidau, Switzerland) on a Leica EM UC7 ultramicrotome (Leica, Wetzlar, Austria) and then examined with a JEM1400 transmission electron microscope (JEOL, Japan). Section preparation and TEM were performed at the Center of Collective Use for Microscopic Analysis of Biological Objects (ICG SB RAS, Novosibirsk, Russia, FWNR-2022-0015). Fixation, dehydration and embedding were performed at the Sector of Structural Cell Biology (ICG SB RAS).

Cell junctions were measured using iTEM software version 5.1 (Olympus, USA) in randomly chosen sections (20 independent measurements per group for each structure per each patient or donor) in a blinded manner by the researcher not involved in data acquisition. Measurements were taken in epithelial cells. For TJ and AJ width and length assessments, 10-20 structures were measured in each group. Three UC patients and two healthy donors were estimated.

#### Data analysis and statistics

A formal power analysis was not conducted to determine sample size in any experiments in this work. Prior to selecting appropriate tests, all datasets were evaluated for normality using the Kolmogorov–Smirnov test. For normally distributed variables, group comparisons were conducted using two-tailed unpaired Student’s t-tests, otherwise the non-parametric Mann–Whitney *U*-test was applied for multiple comparisons, p-values were adjusted using the Benjamini–Hochberg false-discovery rate procedure. In the case of principal component analysis (PCA), group differences were assessed by permutational multivariate analysis of variance (PERMANOVA, adonis function) with 999 permutations, applied to Euclidean distance matrices calculated from Z-score-normalized mass-spectrometry intensities (mean-centered and scaled to unit variance). Statistical significance was defined as p < 0.05 (*), p < 0.01 (**), and p < 0.001 (***). Data visualization was carried out in RStudio 2024.04.2 using the ggplot2 (83)and ggbeeswarm (84) packages.

We used the ARRIVE reporting guideline (85) to draft this manuscript, and the ARRIVE reporting checklist when editing, included in supplementary materials.

## References

1. Di Tommaso N, Gasbarrini A, Ponziani FR. Intestinal barrier in human health and disease. Int J Environ Res Public Health. 2021;18(23):12836.

2. Rescigno M, Chieppa M. Gut-level decisions in peace and war. Vol. 11, Nature Medicine. 2005.

3. Michielan A, D’Incà R. Intestinal Permeability in Inflammatory Bowel Disease: Pathogenesis, Clinical Evaluation, and Therapy of Leaky Gut. Vol. 2015, Mediators of Inflammation. 2015.

4. McDowell C, Farooq U, Haseeb M. Inflammatory Bowel Disease - StatPearls - NCBI Bookshelf. StatPearls Publishing. 2021.

5. DeRoche TC, Xiao SY, Liu X. Histological evaluation in ulcerative colitis. Vol. 2, Gastroenterology Report. 2014.

6. Liverani E, Scaioli E, John Digby R, Bellanova M, Belluzzi A. How to predict clinical relapse in inflammatory bowel disease patients. Vol. 22, World Journal of Gastroenterology. 2016.

7. Villanacci V, Reggiani-Bonetti L, Salviato T, Leoncini G, Cadei M, Albarello L, et al. Histopathology of IBD colitis. A practical approach from the pathologists of the Italian group for the study of the gastrointestinal tract (GIPAD). Vol. 113, Pathologica. 2021.

8. Buda A, Hatem G, Neumann H, D’Incà R, Mescoli C, Piselli P, et al. Confocal laser endomicroscopy for prediction of disease relapse in ulcerative colitis: A pilot study. J Crohns Colitis. 2014;8(4).

9. Chang J, Leong RW, Wasinger VC, Ip M, Yang M, Phan TG. Impaired Intestinal Permeability Contributes to Ongoing Bowel Symptoms in Patients With Inflammatory Bowel Disease and Mucosal Healing. Gastroenterology. 2017;153(3).

10. Geboes K, Riddell R, Öst A, Jensfelt B, Persson T, Löfberg R. A reproducible grading scale for histological assessment of inflammation in ulcerative colitis. Gut. 2000;47(3).

11. Binienda A, Twardowska A, Makaro A, Salaga M. Dietary carbohydrates and lipids in the pathogenesis of leaky gut syndrome: An overview. Int J Mol Sci. 2020;21(21):8368.

12. Vancamelbeke M, Vermeire S. The intestinal barrier: a fundamental role in health and disease. Expert Rev Gastroenterol Hepatol. 2017;11(9):821–34.

13. Belardi B, Hamkins-Indik T, Harris AR, Kim J, Xu K, Fletcher DA. A Weak Link with Actin Organizes Tight Junctions to Control Epithelial Permeability. Dev Cell. 2020;54(6):792–804.

14. Turner JR. Intestinal mucosal barrier function in health and disease. Nat Rev Immunol. 2009;9(11):799–809.

15. Edelblum KL, Turner JR. The tight junction in inflammatory disease: communication breakdown. Curr Opin Pharmacol. 2009;9(6):715–20.

16. Kucharzik T, Walsh S V., Chen J, Parkos CA, Nusrat A. Neutrophil transmigration in inflammatory bowel disease is associated with differential expression of epithelial intercellular junction proteins. American Journal of Pathology. 2001;159(6):2001–9.

17. Mehta S, Nijhuis A, Kumagai T, Lindsay J, Silver A. Defects in the adherens junction complex (E-cadherin/ β-catenin) in inflammatory bowel disease. Cell Tissue Res. 2015;360(3):749–60.

18. Lechuga S, Ivanov AI. Disruption of the epithelial barrier during intestinal inflammation: Quest for new molecules and mechanisms. Biochim Biophys Acta Mol Cell Res. 2017;1864(7):1183–94.

19. Ivanov AI, Parkos CA, Nusrat A. Cytoskeletal regulation of epithelial barrier function during inflammation. American Journal of Pathology. 2010;177(2):512–24.

20. VanDussen KL, Stojmirović A, Li K, Liu TC, Kimes PK, Muegge BD, et al. Abnormal Small Intestinal Epithelial Microvilli in Patients With Crohn’s Disease. Gastroenterology. 2018;155(3):815–28.

21. Goyal N, Rana A, Ahlawat A, Bijjem KR V., Kumar P. Animal models of inflammatory bowel disease: A review. Inflammopharmacology. 2014;22(4):219–33.

22. D. C, Katsikeros R, M. S. Current and Novel Treatments for Ulcerative Colitis. In: Mustafa M. Shennak, editor. Ulcerative Colitis from Genetics to Complications. London: Intech; 2012. p. 189–211.

23. Sluis M Van der, Koning BAE De, Bruijn ACJM De, Velcich A, Meijerink JPP, Goudoever JB Van, et al. Muc2-Deficient Mice Spontaneously Develop Colitis, Indicating That MUC2 Is Critical for Colonic Protection. Gastroenterology [Internet]. 2006 Jul 1 [cited 2021 Oct 9];131(1):117–29. Available from: http://www.gastrojournal.org/article/S0016508506007621/fulltext

24. Ostanin D V., Bao J, Koboziev I, Gray L, Robinson-Jackson SA, Kosloski-Davidson M, et al. T cell transfer model of chronic colitis: Concepts, considerations, and tricks of the trade. Vol. 296, American Journal of Physiology - Gastrointestinal and Liver Physiology. 2009.

25. Borisova MA, Achasova KM, Morozova KN, Andreyeva EN, Litvinova EA, Ogienko AA, et al. Mucin-2 knockout is a model of intercellular junction defects, mitochondrial damage and ATP depletion in the intestinal epithelium. Sci Rep [Internet]. 2020;10(1):21135. Available from: 10.1038/s41598-020-78141-4

26. Walsh S V., Hopkins AM, Chen J, Narumiya S, Parkos CA, Nusrat A. Rho kinase regulates tight junction function and is necessary for tight junction assembly in polarized intestinal epithelia. Gastroenterology. 2001;121(3).

27. Hwang JS, Vo TTL, Kim M, Cha EH, Mun KC, Ha E, et al. Involvement of RhoA/ROCK Signaling Pathway in Methamphetamine-Induced Blood-Brain Barrier Disruption. Biomolecules. 2025;15(3).

28. Eichele DD, Kharbanda KK. Dextran sodium sulfate colitis murine model: An indispensable tool for advancing our understanding of inflammatory bowel diseases pathogenesis. Vol. 23, World Journal of Gastroenterology. 2017.

29. Eden K. Adoptive transfer colitis. In: Methods in Molecular Biology. 2019.

30. Boldyreva L V., Morozova M V., Saydakova SS, Kozhevnikova EN. Fat of the gut: Epithelial phospholipids in inflammatory bowel diseases. Vol. 22, International Journal of Molecular Sciences. MDPI; 2021. p. 11682.

31. Vetrano S, Rescigno M, Rosaria Cera M, Correale C, Rumio C, Doni A, et al. Unique Role of Junctional Adhesion Molecule-A in Maintaining Mucosal Homeostasis in Inflammatory Bowel Disease. Gastroenterology. 2008;135(1):173–84.

32. Kuo WT, Zuo L, Odenwald MA, Madha S, Singh G, Gurniak CB, et al. The Tight Junction Protein ZO-1 Is Dispensable for Barrier Function but Critical for Effective Mucosal Repair. Gastroenterology. 2021;161(6):1924–39.

33. Oshima T, Miwa H, Joh T. Changes in the expression of claudins in active ulcerative colitis. Journal of Gastroenterology and Hepatology (Australia). 2008;23:146–50.

34. Luettig J, Rosenthal R, Barmeyer C, Schulzke JD. Claudin-2 as a mediator of leaky gut barrier during intestinal inflammation. Tissue Barriers. 2015;3(1–2):e977176.

35. Wang K, Ding Y, Xu C, Hao M, Li H, Ding L. Cldn-7 deficiency promotes experimental colitis and associated carcinogenesis by regulating intestinal epithelial integrity. Oncoimmunology. 2021;10(1):1923910.

36. Robinson SC, Chaudhary R, Jiménez-Saiz R, Rayner LGA, Bayer L, Jordana M, et al. Kaiso-induced intestinal inflammation is preceded by diminished E-cadherin expression and intestinal integrity. PLoS One. 2019;14(6):e0217220.

37. Ohta H, Sunden Y, Yokoyama N, Osuga T, Lim SY, Tamura Y, et al. Expression of apical junction complex proteins in duodenal mucosa of dogs with inflammatory bowel disease. Am J Vet Res. 2014;75(8):746–51.

38. Ivanov AI, Nusrat A, Parkos CA. The epithelium in inflammatory bowel disease: Potential role of endocytosis of junctional proteins in barrier disruption. In: Novartis Foundation Symposium 263. Chichester, UK: John Wiley & Sons; 2004. p. 115–32.

39. Xing T, Benderman LJ, Sabu S, Parker J, Yang J, Lu Q, et al. Tight Junction Protein Claudin-7 Is Essential for Intestinal Epithelial Stem Cell Self-Renewal and Differentiation. CMGH. 2020;9(4):641–59.

40. Fredericks E, Dealtry G, Roux S. β -Catenin Regulation in Sporadic Colorectal Carcinogenesis: Not as Simple as APC. Can J Gastroenterol Hepatol. 2018;2018(1):4379673.

41. Bruewer M, Utech M, Ivanov AI, Hopkins AM, Parkos CA, Nusrat A. Interferon-γ induces internalization of epithelial tight junction proteins via a macropinocytosis-like process. The FASEB Journal. 2005;19(8):923–33.

42. Bruewer M, Luegering A, Kucharzik T, Parkos CA, Madara JL, Hopkins AM, et al. Proinflammatory Cytokines Disrupt Epithelial Barrier Function by Apoptosis-Independent Mechanisms. The Journal of Immunology. 2003;171(11):6164–72.

43. Musch MW, Walsh-Reitz MM, Chang EB. Roles of ZO-1, occludin, and actin in oxidant-induced barrier disruption. Am J Physiol Gastrointest Liver Physiol. 2006;290(2):222–31.

44. Oshitani N, Watanabe K, Nakamura S, Fujiwara Y, Higuchi K, Arakawa T. Dislocation of tight junction proteins without F-actin disruption in inactive Crohn’s disease. Int J Mol Med. 2005;15(3):407–10.

45. Troyanovsky RB, Indra I, Troyanovsky SM. Actin-dependent α-catenin oligomerization contributes to adherens junction assembly. Nature Communications. 2025;16(1).

46. Song H, Zhang J, He W, Wang P, Wang F. Activation of Cofilin Increases Intestinal Permeability via Depolymerization of F-Actin During Hypoxia in vitro. Front Physiol. 2019;10:1455.

47. van Goor D, Hyland C, Schaefer AW, Forscher P. The role of actin turnover in retrograde actin network flow in neuronal growth cones. PLoS One. 2012;7(2).

48. Sakata A, Ochiai T, Shimeno H, Hikishima S, Yokomatsu T, Shibuya S, et al. Acid sphingomyelinase inhibition suppresses lipopolysaccharide-mediated release of inflammatory cytokines from macrophages and protects against disease pathology in dextran sulphate sodium-induced colitis in mice. Immunology. 2007;122(1):54–64.

49. Bauer J, Liebisch G, Hofmann C, Huy C, Schmitz G, Obermeier F, et al. Lipid alterations in experimental murine colitis: Role of ceramide and imipramine for matrix metalloproteinase-1 expression. PLoS One. 2009;4(9):e7197.

50. Shores DR, Binion DG, Freeman BA, Baker PRS. New insights into the role of fatty acids in the pathogenesis and resolution of inflammatory bowel disease. Inflamm Bowel Dis. 2011;17(10):2192–204.

51. Suh JH, Saba JD. Sphingosine-1-phosphate in inflammatory bowel disease and colitis-associated colon cancer: The fat’s in the fire. Transl Cancer Res. 2015;4(5):469.

52. Radner FPW, Marrakchi S, Kirchmeier P, Kim GJ, Ribierre F, Kamoun B, et al. Mutations in CERS3 Cause Autosomal Recessive Congenital Ichthyosis in Humans. PLoS Genet. 2013;9(6).

53. Levy M, Futerman AH. Mammalian ceramide synthases. IUBMB Life. 2010;62(5):347–56.

54. Diab J, Hansen T, Goll R, Stenlund H, Ahnlund M, Jensen E, et al. Lipidomics in Ulcerative Colitis Reveal Alteration in Mucosal Lipid Composition Associated with the Disease State. Inflamm Bowel Dis. 2019;25(11):1780–7.

55. Bazarganipour S, Hausmann J, Oertel S, El-Hindi K, Brachtendorf S, Blumenstein I, et al. The lipid status in patients with ulcerative colitis: Sphingolipids are disease-dependent regulated. J Clin Med. 2019;8(7):971.

56. Kim YR, Volpert G, Shin KO, Kim SY, Shin SH, Lee Y, et al. Ablation of ceramide synthase 2 exacerbates dextran sodium sulphate-induced colitis in mice due to increased intestinal permeability. J Cell Mol Med. 2017;21(12):3565–78.

57. Spiegel S, Foster D, Kolesnick R. Signal transduction through lipid second messengers. Curr Opin Cell Biol. 1996;8(2):159–67.

58. Liu X, Shu S, Billington N, Williamson CD, Yu S, Brzeska H, et al. Mammalian nonmuscle myosin II binds to anionic phospholipids with concomitant dissociation of the regulatory light chain. Journal of Biological Chemistry. 2016;291(48):24828–37.

59. Bieberich E. Ceramide signaling in cancer and stem cells. Future Lipidol. 2008;3(3):273–300.

60. Krishnamurthy K, Wang G, Silva J, Condie BG, Bieberich E. Ceramide regulates atypical PKCζ/λ-mediated cell polarity in primitive ectoderm cells: A novel function of sphingolipids in morphogenesis. Journal of Biological Chemistry. 2007;282(5):3379–90.

61. Bock J, Liebisch G, Schweimer J, Schmitz G, Rogler G. Exogenous sphingomyelinase causes impaired intestinal epithelial barrier function. World J Gastroenterol. 2007;13(39):5217–25.

62. Lee DBN, Jamgotchian N, Allen SG, Abeles MB, Ward HJ. A lipid-protein hybrid model for tight junction. Am J Physiol Renal Physiol. 2008;295(6):1601–12.

63. McGuinn KP, Mahoney MG. Lipid rafts and detergent-resistant membranes in epithelial keratinocytes. Methods in Molecular Biology. 2014;1195:133–44.

64. Saslowsky DE, Thiagarajah JR, McCormick BA, Lee JC, Lencer WI. Microbial sphingomyelinase induces RhoA-mediated reorganization of the apical brush border membrane and is protective against invasion. Mol Biol Cell. 2016;27(7):1120–30.

65. Kwik J, Boyle S, Fooksman D, Margolis L, Sheetz MP, Edidin M. Membrane cholesterol, lateral mobility, and the phosphatidylinositol 4,5-bisphosphate-dependent organization of cell actin. Proc Natl Acad Sci U S A. 2003;100(24):13964–9.

66. Danielsen EM, Hansen GH. Lipid rafts in epithelial brush borders: Atypical membrane microdomains with specialized functions. Biochim Biophys Acta Biomembr. 2003;1617(1–2):1–9.

67. Oliferenko S, Paiha K, Harder T, Gerke V, Schwärzler C, Schwarz H, et al. Analysis of CD44-containing lipid rafts: Recruitment of annexin II and stabilization by the actin cytoskeleton. Journal of Cell Biology. 1999;146(4):843–54.

68. Bowie R V., Donatello S, Lyes C, Owens MB, Babina IS, Hudson L, et al. Lipid rafts are disrupted in mildly inflamed intestinal microenvironments without overt disruption of the epithelial barrier. Am J Physiol Gastrointest Liver Physiol. 2012;302(8):781–93.

69. Schmelz EM, Roberts PC, Kustin EM, Lemonnier LA, Sullards MC, Dillehay DL, et al. Modulation of intracellular β-catenin localization and intestinal tumorigenesis in vivo and in vitro by sphingolipids. Cancer Res. 2001;61(18):6723–9.

70. Li Y, Nicholson RJ, Summers SA. Ceramide signaling in the gut. Mol Cell Endocrinol. 2022;544:111554.

71. Günzel D, Yu ASL. Claudins and the modulation of tight junction permeability. Physiol Rev. 2013;93(2):525.

72. Krause G, Winkler L, Mueller SL, Haseloff RF, Piontek J, Blasig IE. Structure and function of claudins. Biochim Biophys Acta Biomembr. 2008;1778(3):631–45.

73. Capaldo CT. Claudin Barriers on the Brink: How Conflicting Tissue and Cellular Priorities Drive IBD Pathogenesis. Int J Mol Sci. 2023;24(10):8562.

74. Chen J, Wei Z, Wang Y, Long M, Wu W, Kuca K. Fumonisin B1: Mechanisms of toxicity and biological detoxification progress in animals. Food and Chemical Toxicology. 2021;149:111977.

75. Li M, Liu S, Tan L, Luo Y, Gao Z, Liu J, et al. Fumonisin B1 induced intestinal epithelial barrier damage through endoplasmic reticulum stress triggered by the ceramide synthase 2 depletion. Food and Chemical Toxicology. 2022;166:113263.

76. Li X, Cao C, Zhu X, Li X, Wang K. Fumonisins B1 exposure triggers intestinal tract injury via activating nuclear xenobiotic receptors and attracting inflammation response. Environmental Pollution. 2020;267:115461.

77. Bouhet S, Hourcade E, Loiseau N, Fikry A, Martinez S, Roselli M, et al. The mycotoxin fumonisin B1 alters the proliferation and the barrier function of porcine intestinal epithelial cells. Toxicological Sciences. 2004;77(1):165–71.

78. Kanai T, Kawamura T, Dohi T, Makita S, Nemoto Y, Totsuka T, et al. T H1/T H2-mediated colitis induced by adoptive transfer of CD4 +CD45RB high T lymphocytes into nude mice. Inflamm Bowel Dis. 2006;12(2).

79. Bradford MM. A rapid and sensitive method for the quantitation of microgram quantities of protein utilizing the principle of protein-dye binding. Anal Biochem. 1976;72(1–2):248–54.

80. Pang Z, Chong J, Zhou G, De Lima Morais DA, Chang L, Barrette M, et al. MetaboAnalyst 5.0: Narrowing the gap between raw spectra and functional insights. Nucleic Acids Res. 2021;49(1):388–96.

81. Xu S, Hu E, Cai Y, Xie Z, Luo X, Zhan L, et al. Using clusterProfiler to characterize multiomics data. Nat Protoc. 2024;19(11).

82. Carlson Marc. Bioconductor. [cited 2026 Jan 15]. org.Mm.eg.db. Genome wide annotation for Mouse. Available from: https://bioconductor.org/packages/release/data/annotation/html/org.Mm.eg.db.html

83. Wickham H. ggplot2 - Elegant Graphics for Data Analysis | Hadley Wickham | Springer. Springer Science & Business Media. 2017.

84. Clarke Erik, Sherrill-Mix Scott, Dawson Charlotte. ggbeeswarm: Categorical Scatter (Violin Point) Plots [Internet]. [cited 2026 Jan 15]. Available from: https://cran.r-project.org/web/packages/ggbeeswarm/

85. du Sert NP, Hurst V, Ahluwalia A, Alam S, Avey MT, Baker M, et al. The arrive guidelines 2.0: Updated guidelines for reporting animal research. PLoS Biol. 2020;18(7).

